# Kinetic principles underlying pioneer function of GAGA transcription factor in live cells

**DOI:** 10.1101/2021.10.21.465351

**Authors:** Xiaona Tang, Taibo Li, Sheng Liu, Jan Wisniewski, Qinsi Zheng, Yikang Rong, Luke D. Lavis, Carl Wu

## Abstract

How pioneer factors interface with chromatin to promote accessibility for transcription control is poorly understood in vivo. Here, we directly visualize chromatin association by the prototypical GAGA pioneer factor (GAF) in live *Drosophila* hemocytes. Single-particle tracking reveals that the majority of GAF is chromatin-bound, with a stable-binding fraction showing nucleosome-like confinement residing on chromatin for over 2 minutes, far longer than the dynamic range of most transcription factors. These kinetic properties require the full complement of GAF’s DNA-binding, multimerization and intrinsically disordered domains, and are autonomous from recruited chromatin remodelers NURF and PBAP, whose activities primarily benefit GAF’s neighbors such as HSF. Evaluation of GAF kinetics together with its endogenous abundance indicates that despite on-off dynamics, GAF constitutively and fully occupies chromatin targets, thereby providing a temporal mechanism that sustains open chromatin for transcriptional responses to homeostatic, environmental, and developmental signals.

## Main

*Drosophila* GAGA factor (GAF), a ubiquitous and essential Zn finger transcription factor (TF) encoded by the *Trithorax-like* (*Trl*) gene, is a multimeric protein complex that binds specifically to clusters of adjacent GAGAG sequences on numerous genes, including homeotic, steroid- and heat shock-response genes^1–4^. GAF regulates transcription by interactions with the TAF3 and TAF4 components of the TFIID general transcription factor^5–7^, the NELF elongation factor^8^, and antagonism to histone H1-mediated transcriptional repression^9^. In addition, as a pioneer transcription factor^10,11^, GAF is capable of binding to reconstituted nucleosomes^12^, directly recruiting chromatin remodelers NURF, PBAP and other factors^6,13–15^ to create accessible chromatin for neighboring non-pioneer factors^16^ and assembly of the paused RNA Polymerase II (Pol II)^17,18^. Beneficiaries of GAF pioneering activity in *Drosophila* include - heat shock factor HSF^19^, coactivator CBP^19,20^, Polycomb repressor PHO (Pleiohomeotic)^21^, and the insulator binding complex LBC (Large Boundary Complex)^22^.

Genome-wide analysis reveals that GAF is enriched *in vivo* at promoters as well as distal *cis*-regulatory regions comprising several thousand targets which often include clusters of tandem GA repeats^17,23,24^. GAF-specific RNA or protein depletion experiments have importantly demonstrated the in vivo function of GAF in generating chromatin accessibility in the *Drosophila* embryo and in cultured cells^17,18,24,25^. Despite these advances over decades of research, unifying principles for pioneering activity of TFs such as GAF have remained elusive. How pioneer factors differ from other sequence-specific transcription factors and the kinetic mechanisms by which pioneers perform their key genomic functions are subjects of continuing debate.

To elucidate the underlying mechanisms of this pleiotropic transcription factor, we have studied the kinetics of GAF diffusion in the nucleoplasm and on genomic chromatin by single-particle tracking (SPT) in live *Drosophila* hemocytes under different genetic contexts. In comparison, we also measured the kinetic parameters for HSF at normal and heat-stressed conditions. We then determined GAF and HSF protein levels in hemocytes, curated existing databases for numbers of genomic targets and integrated these parameters with the measured kinetics to obtain the target occupancy of each factor in vivo. Our findings uncover crucial quantitative principles for pioneering and maintaining chromatin accessibility over extended time periods even when the responsible factors continuously bind to and dissociate from their chromatin targets in a dynamic manner.

## Results

### GAF imaging quantifies major chromatin binding fractions and prolonged residence times

*Drosophila Trl* encodes two isoforms of GAF that harbor the N-terminal POZ/BTB domain (hereafter called POZ), the central zinc-finger (ZF)-containing DNA binding domain (DBD), and long and short C-terminal Q-rich domains **(Fig. 1A)^4,26^**. We used CRISPR-Cas9-based gene editing to insert a HaloTag at the N-terminus of endogenous *Trl* **(Fig. 1A** and **Fig. S1A-D)**. The Halo knock-in strain is homozygous-viable, expressing long and short Halo-GAF isoforms (GAF^L^ and GAF^S^) as the sole source. Halo-GAF binds to numerous loci on salivary gland polytene chromosomes, consistent with immunostaining studies^12,26,27^ **(Fig. 1B)**, and appears in multiple nuclear foci in diploid circulating larval hemocytes (>90% plasmatocytes, counterpart of mammalian macrophages^28^), similar to nuclear foci in the S2 cell line^29^ **(Fig. 1C)**. We also constructed transgenic flies expressing C-terminal tagged isoforms GAF^L^-Halo and GAF^S^-Halo, which exhibit tissue-specific expression^26^ **(Fig. S1F)** and are functionally active, as indicated by rescue of *Trl^13C^/Trl^R67^* lethal alleles^3^. Together, the data demonstrate that fusion of HaloTag does not interfere with the localization and essential functions of GAF.

**Figure 1.**
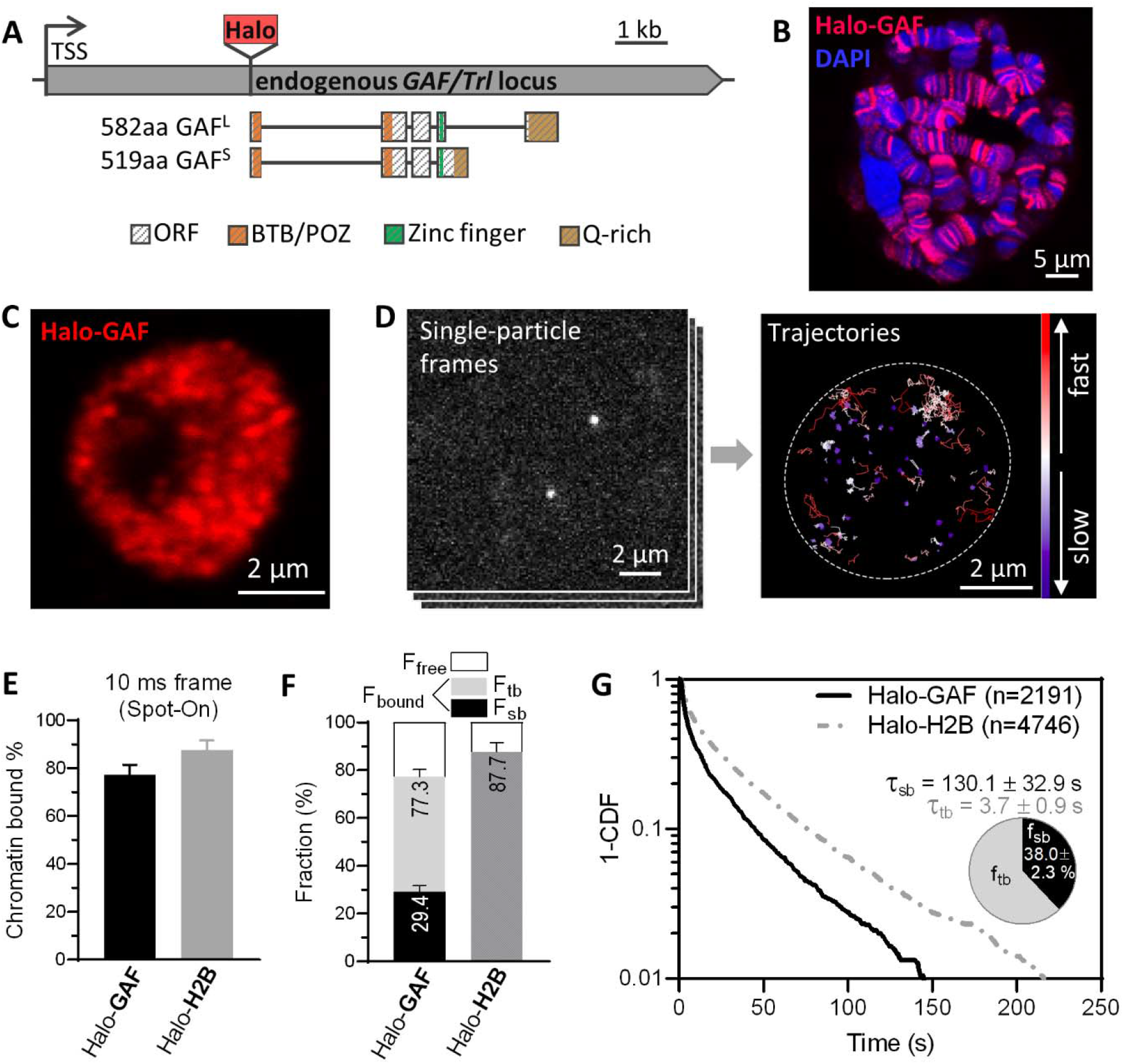
Chromatin-binding dynamics of GAF shown by single-particle tracking (SPT) in live *Drosophila* hemocytes. (A) Diagram of HaloTag (Halo) knock-in at the N-terminus of endogenous *GAF/Trl* locus. Protein domains in color. TSS, transcription start site. C-terminal GAF-HaloTag fusions for two splicing isoforms GAF^S^ (short) and GAF^L^ (long) are shown in Fig S1. (B) Halo-GAF binds to specific polytene chromosome loci in fixed 3^rd^ instar larval salivary gland nuclei, revealed by JF552 fluorescence (red); DNA counterstained with DAPI (blue). (C) Confocal distribution of Halo-GAF (JF552) foci in fixed 3rd instar larval hemocyte nuclei (predominantly diploid plasmatocytes in G2 phase of the cell cycle). (D) SMT frames and superimposed trajectories of Halo-GAF in live hemocytes, color-coded according to diffusion coefficients. Dashed oval marks the nucleus. (E) Spot-On kinetic modeling of fast-tracking data shows chromatin-bound and free fractions for Halo-GAF and Halo-H2B. Results are mean ± SD from three biological replicates. (F) Chromatin-free fraction (F_free_), global stable- and transient-binding fractions (F_sb_ and F_tb_) of Halo-GAF extracted from fast- and slow-tracking data in (E) and (G), with error propagation. (G) Survival probability curves (1-CDF) plotted from apparent dwell times of thousands (n) of single-particle chromatin-binding events for Halo-GAF. Average residence times for stable- (τ_sb_) and transient- (τ_tb_) binding by Halo-GAF are corrected by using Halo-H2B as a ‘non-dissociating’ standard. Pie charts show stable-(f_sb_) and transient-binding (f_tb_) fractions. Mean and SD provided. Errors represent bootstrapped SD.

We investigated the live-cell dynamics of tagged GAF species in live hemocytes **(Fig. S2A)** by SPT, using a ‘fast-tracking’ regime (10 ms/frame) in dSTORM mode^30–32^ to measure slow- and fast-diffusing molecules **(Fig. 1D and Movie S1)**, and quantified diffusion coefficients with a robust, displacement-based, analytical protocol (Spot-On)^33^ **(Fig. 1E** and **Fig. S2B)**. All Halo-tagged GAF versions display similar slow and fast diffusivities that are within similar dynamic range demonstrated by histone H2B **(Fig. 1E** and **Fig. S2C)**, with >75% and >85% slow-diffusing fraction (chromatin-bound, F_bound_), respectively. The diffusion coefficient (D) of the bound fraction (D_bound_ 0.004-0.005 um^2^/s) is two orders of magnitude lower than the free fraction (D_free_ 0.75-0.8 um^2^/s) **(Fig. S2E-F)**.

A ‘slow-tracking’ regime (500 ms/frame) to motion-blur fast particles allowed selective detection of long- and short-lived chromatin-bound populations **(Fig. S3A-C** and **Movie S2)**. We calculated 1-CDF of dwell times to generate survival curves demonstrating apparent GAF dissociation over time. Halo-GAF and GAF^L^-Halo show similar profiles, with slightly faster decay for the GAF^S^-Halo isoform **(Fig. S3D)**. Fitting to a double exponential function **(Fig. S3E)** enabled calculation of long- and short-lived (called stable and transient) binding fractions, f_sb_ and f_tb_, and average stable and transient residence times, τ_sb_ and τ_tb_, after correction for photobleaching and out-of-focus chromatin motions using the apparent dissociation of Halo-H2B as an ‘nondissociative’ standard within the experimental timescale of several minutes **(Fig. S3D)**. The stable binding fraction f_sb_ multiplied by total binding fraction F_bound_ gives the overall stable-binding fraction F_sb_= 30-35 % **(Fig. 1F** and **S2D)**. All stable-binding GAF fractions display strikingly protracted residence times τ_sb_: 130 s for Halo-GAF, 141 s for GAF^L^-Halo, and 85 s for GAF^S^-Halo, which is 20- to 30-fold longer than the transient residence time τ_tb_ of ^~^4 s **(Fig. 1G** and **Fig. S3F)**, and longer than τ_sb_ values measured for many mammalian transcription factors ^32,34–38^. stable- and transient-binding are assumed to occur at cognate and non-specific sites, respectively.

### POZ, Q-rich, and DBD domains of GAF contribute to stable chromatin binding

Purified, bacterially expressed GAF associates in oligomeric complexes ranging from monomers to multimers^39,40^, and GAF complexes are also observed in nuclear extracts of S2 cells. Multimerization is mediated by the POZ and Q-rich domains of GAF^39–41^. To investigate how GAF domains contribute to particle dynamics, we used CRISPR-Cas9 to engineer Halo-GAF deletions in the POZ domain (*ΔPOZ*), the zinc finger (*ZF^9^; ZF^10^*), and the long and short Q-rich domains (*ΔQ*) **(Fig. 2A** and **Fig. S4A-E)**.

**Figure 2.**
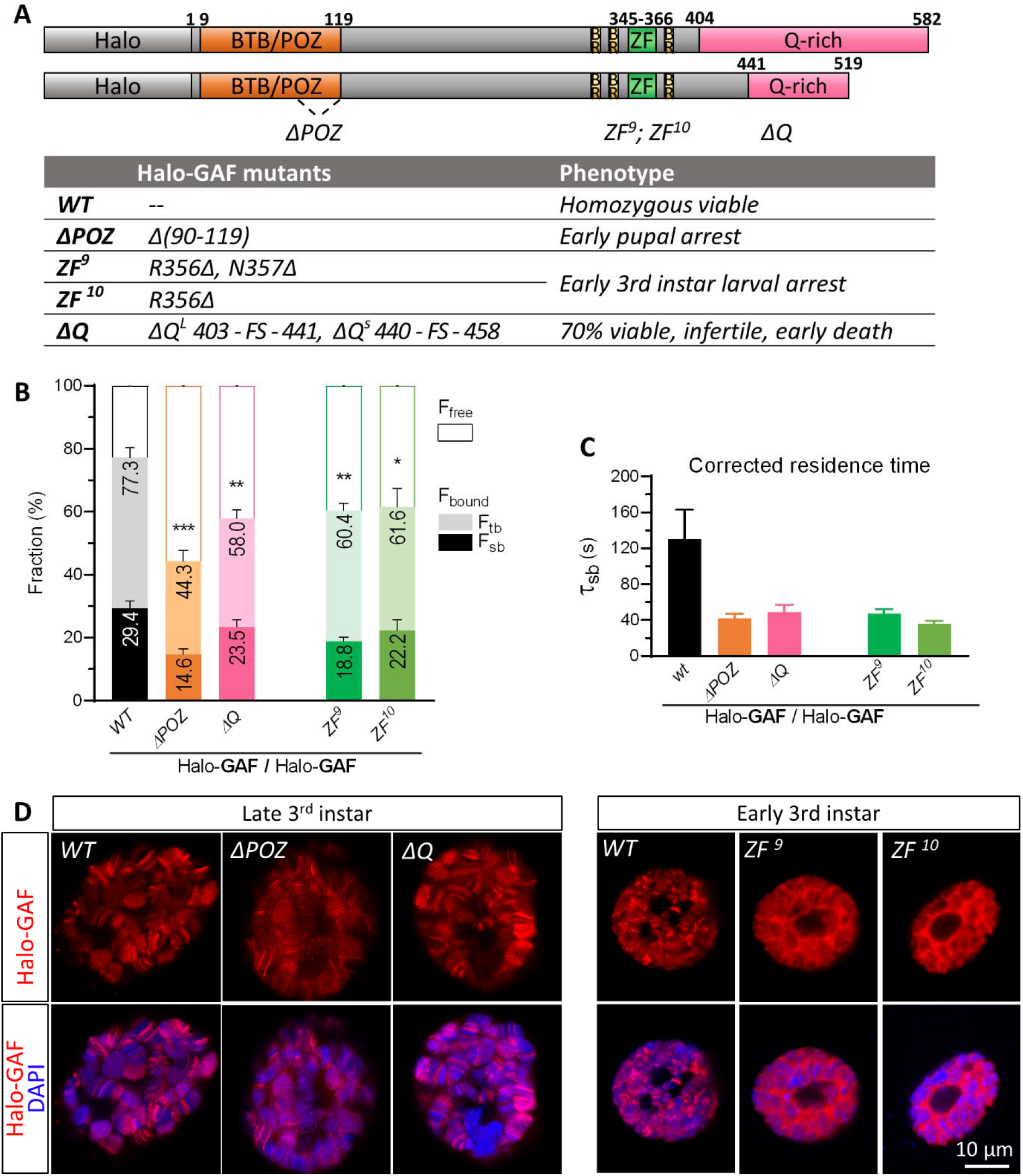
POZ, Q-rich, and DBD domains of GAF all contribute to stable chromatin binding and long residence times. (A) Schematics of Halo-GAF^L^ (long) and Halo-GAF^S^ (short) isoforms with functional domains to scale. BTB/POZ: Broad-complex, Tramtrack and Bric-à-brac/Poxvirus and Zinc finger, BR: basic region, ZF: zinc finger, Q-rich: glutamine-rich. Halo-GAF deletions were generated by CRISPR-Cas9. *ΔPOZ* contains a 90bp deletion in the second exon, generating a 30-AA deletion (*Δ90-119*) of the POZ domain, which includes G93 and L112 that are essential for transcription activation^29^. The deleted Arg and Asn amino acids in the zinc finger (*ZF^9^, ZF^10^*) make contact with ‘GAG’ of the consensus GAF binding site(Omichinski et al. 1997). *ΔQ* contains small deletions at the beginning of Q-rich domains of both long and short isoforms, resulting in frame shift and truncation of Q-rich domains. Table reports mutant phenotypes. FS: frameshift. (B) Global chromatin-bound and free fractions (%) for wild type (*WT*) and mutant Halo-GAF. Fast and slow-tracking results with propagated errors. *, *p*< 0.05; **, *p* < 0.01; ***, *p* < 0.001, unpaired *t*-test for fast-tracking (n=3-4). (C) Average residence times for *WT* and mutant Halo-GAF, corrected as in Fig. 1. Error bars represent bootstrapped SD. (D) Halo-GAF distribution on fixed salivary gland (SG) polytene nuclei for *WT* and GAF mutants. *ΔPOZ* and *ΔQ* nuclei are from late 3rd instar larvae with *WT* nuclei from the same stage as control. *ZF^9^* and *ZF^10^* polytene nuclei are from early instar larvae with *WT* nuclei from the same stage as control. Red: Halo-GAF; blue: DAPI. One representative confocal z-section is shown.

Homozygous *ΔPOZ* and *ZF^9^* or *ZF^10^* mutants arrest at early pupal and early 3^rd^ instar larval stages, respectively, while homozygous *ΔQ* is 70% viable, producing infertile adults with impaired longevity **(Fig. 2A)**. These manifest phenotypes, although late in development owing to perdurance of the untagged WT GAF contributed by the heterozygous mother to homozygous oocytes, indicate that all three GAF domains are essential for *Drosophila* viability.

SPT using fast and slow tracking regimes for Halo-GAF *ΔPOZ, ZF^9^, ZF^10^* and *ΔQ* mutants in 3^rd^ instar larval hemocytes found that all three domain mutants display substantial reductions in F_sb_ and τ_sb_ for the slow-diffusing fraction **(Fig. 2B-C** and **Fig. S5A-C)**. Disruption of the POZ domain reduces F_sb_ from 29% to 15%, and τ_sb_ from 130 s to 42 s, demonstrating the important contribution of this multimerization domain to stable chromatin association. Deletion of Q-rich domains shows a modest reduction of F_sb_ from 29% to 24%, and reduces the τ_sb_ from 130 s to 49 s. Interestingly, under fast-tracking, *ΔPOZ* and *ΔQ* proteins exhibit similar D_free_ but larger D_bound_ compared to *WT* GAF, suggesting a more diffusive binding mode **(Fig. S5D-E)**. These results are consistent with the specific binding patterns of *ΔPOZ* and *ΔQ* on fixed polytene chromosomes (Fig. 2D)

The zinc finger mutants *ZF^9^* and *ZF^10^* also show reductions in F_sb_ to 19% and 22%, and τ_sb_ to ^~^40 s in hemocytes **(Fig. 2B-C and Fig. S5A-C**), similar to *ΔPOZ* and *ΔQ*. However, unlike *ΔPOZ* and *ΔQ*, both *ZF^9^* and *ZF^10^* mutants exhibit loss of the specific binding pattern on polytene chromosomes and increase of nucleoplasmic distribution **(Fig. 2D)**. These results confirm the crucial function of the DNA-binding zinc finger for site-specific chromatin binding, and report aberrant, mechanistically unclear diffusive behavior of *ZF^9^* and *ZF^10^* mutant proteins. Notably, the time-averaged Mean Squared Displacement (MSD) curves from slow-tracking of *ZF^9^* and *ZF^10^* show an initial steep rise followed by a plateau after 10 s, a profile dramatically different from *WT* GAF, *ΔPOZ, ΔQ*, and H2B HaloTag fusions, all of which display a linear or Brownian increase of the MSD over a 60 s timescale **(Fig. S5F)**.

### Chromatin binding kinetics by GAF is independent of recruited remodelers NURF and PBAP

At cognate GAGAG sites on *Drosophila* chromatin, GAF recruits NURF and PBAP, ATP-dependent chromatin remodelers of the ISWI and SWI/SNF families, respectively, to drive DNA accessibility for neighboring transcription factors and establish promoter-proximal paused Pol II^13,17,19,24,42,43^. This process begins during *Drosophila* embryogenesis, when GAF and the Zelda pioneer factor are individually required to activate and remodel the chromatin accessibility landscape for widespread zygotic transcription^25,29^. However, it was unclear whether the recruitment of NURF and PBAP by GAF is required to assist its own chromatin binding as a pioneer factor. To address this, we performed SPT of GAF^L^-Halo and GAF^S^-Halo on 3^rd^ instar larval hemocytes isolated from *bap170 and nurf301/E*(*bx*) mutants for unique subunits in PBAP^43,44^ and NURF^19,43^ complexes, respectively **(Fig. 3A)**. The results show little or small changes in the F_bound_, D_bound_ and τ_sb_ values of the mutants **(Fig. 3B-C** and **Fig. S6A-F)**. GAF^L^-Halo and GAF^S^-Halo isoforms also show no qualitative global binding changes on polytene chromosomes in the *bap170* and *nurf301* mutants, although changes in a minority of chromosomal loci might escape detection **(Fig. 3D)**. (By contrast, changes of HSF-Halo binding can be detected in mutants; see below). These findings are generally consistent with ChIP-Seq studies showing similar average GAF binding genome-wide in PBAP-depleted S2 cells **(Fig. S7)**, although partial GAF binding is observed in a subset of *Drosophila* promoters (685 promoters displaying reductions in paused RNA Pol II and chromatin accessibility)^24^. Taken together, our live-cell SPT results indicate that GAF chromatin binding and dwell time are largely autonomous from NURF and PBAP, although other remodeling activities are not excluded^45^. We conclude that the main beneficiaries of NURF and PBAF recruitment and nucleosome remodeling activities are GAF’s ‘non-pioneer’ neighbors.

**Figure 3.**
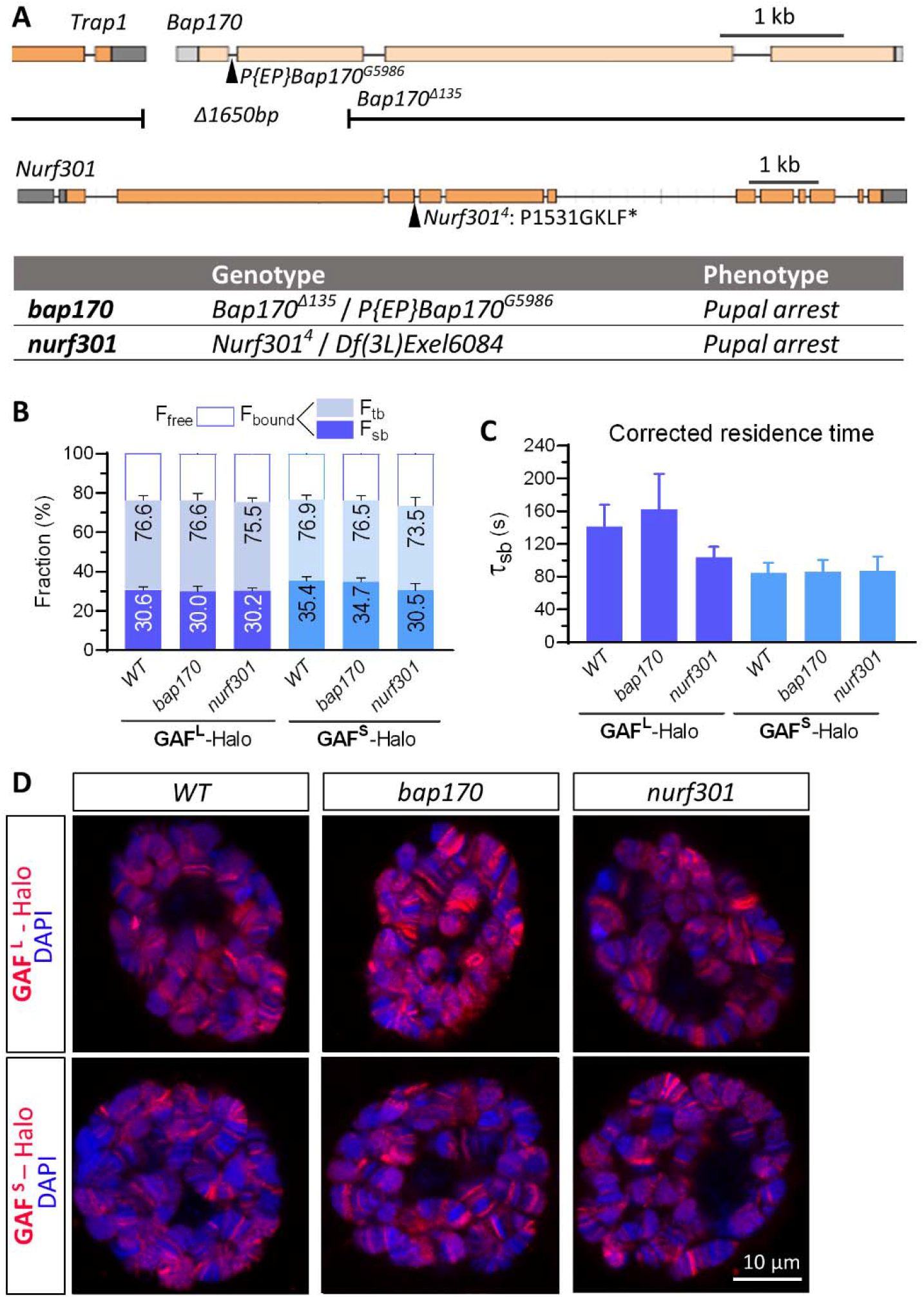
Chromatin binding by GAF is largely independent of remodelers PBAP and NURF. (A) *Bap170* and *Nurf301/E*(*bx*) mutants and phenotypes. *Bap170^Δ135^* harbors a 1650bp deletion spanning the promoter, 5’UTR and first two exons of *Bap170, P{EP}Bap170^G5986^* has a *P{EP}* element insertion at 447 bp downstream from the transcription start site in the first intron of the gene. *Bap170^Δ135^* abolishes *Bap170* expression and strongly reduces polybromo protein level at larval stage, causing pupal lethality when homozygous(Carrera et al. 2008), while flies homozygous for the *P{EP}Bap170^G5986^* die primarily as pharate adults, with only 3% male homozygous adults(Chalkley et al. 2008). Transheterozygous *Bap170^Δ135^/P{EP}Bap170^G5986^* (denoted as *bap170*) also exhibits pupal arrest. *Nurf301^4^* contains a splice-donor site mutation that blocks splicing of the fourth intron. The aberrant transcript introduces four additional amino acids (GKLF) and an in-frame stop codon after P1531, truncating C-terminal 609-1230 aa including the essential PHD finger and Bromodomain(Badenhorst et al. 2002). *Nurf301^4^* is pupal-lethal with <60% pupariation rate(Badenhorst et al. 2005). When transheterozygous with a deletion allele spanning 31 genes including *Nurf301, Nurf301^4^/Df(3L)Exel6084* (denoted as *nurf301*) also shows pupal arrest. (B) Global chromatin-bound fractions for GAF^L^-Halo and GAF^S^-Halo in wild type (*WT*), *bap170* and *nurf301* mutants. Fast and slow tracking results with propagated errors. All three strains express transgenic GAF^L^-Halo or GAF^S^-Halo under natural regulation, in the presence of endogenous, untagged GAF. See methods for genotypes of *WT, bap170*, and *nurf301*. (C) Corrected average residence times for GAF^L^-Halo and GAF^S^-Halo in *WT, bap170* and *nurf301* mutants. Error bars represent bootstrapped SD. (D) GAF^L^-Halo and GAF^S^-Halo distributions on fixed salivary gland polytene nuclei for *WT, bap170* and *nurf301* mutants.

### Heat shock increases chromatin-binding fraction of HSF without affecting dwell time

The recruitment of chromatin remodelers by GAF increases accessibility to facilitate binding of Heat Shock Factor (HSF) to the tripartite Heat Shock Element (HSE) adjacent to GAF binding sites at heat shock promoters^18,46–49^. Under normal conditions, HSF is predominantly monomeric, with low (submicromolar) affinity of its winged-helix DBD for an NGAAN sequence^50,51^; heat shock induces HSF trimerization and juxtaposition of three DBDs for high-affinity binding to HSEs containing triple NGAAN sequences in alternating orientation^52–57^.

To measure HSF dynamics, we constructed a transgenic HSF-Halo strain under natural expression control, and verified the functionality of HSF-Halo by rescue of *P{PZ}Hsf^03091^/Hsf^3^* lethal alleles^58,59^. We further validated HSF-Halo functions by confocal imaging of fixed polytene nuclei, which showed that HSF-Halo is mostly nucleoplasmic at room temperature (RT), except for low binding to few sites including *Hsp83* harboring very high affinity HSEs; heat shock at 37.5 °C for 10 or 30 min induced strong HSF binding to many more chromosomal loci, most prominently reported at *Hsp* genes^52,60^ **(Fig. S8A)**. This inducible pattern of HSF binding on heat shock is partially reduced in mutants for *Trl, Bap170* and *Nurf301* **(Fig. S8A)**. (Note that there are GAF-independent HSF targets in the genome^18^, for which changes of HSF binding at corresponding polytene loci would not be expected). Imaging of fixed hemocytes shows that the heterogeneous distribution of HSF-Halo changes on heat shock to a more punctate pattern including several prominent foci, consistent with previous studies **(Fig. 4A**)^52,61,62^.

**Figure 4.**
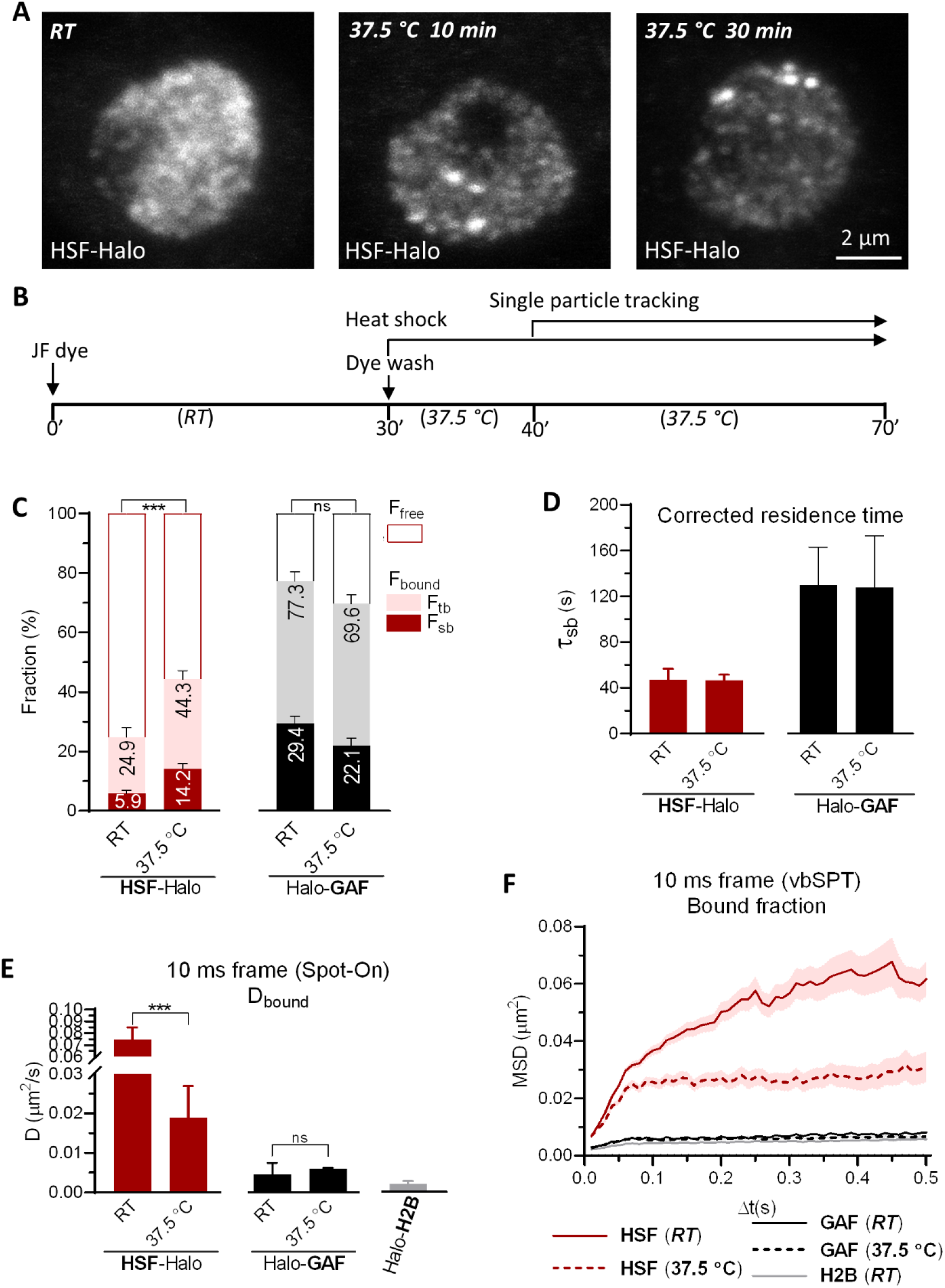
Heat shock increases chromatin binding fraction of HSF without affecting dwell time. (A) Maximum-intensity z-stack projection of HSF-Halo in fixed hemocytes. HSF-Halo forms several prominent foci upon heat shock at 37.5°C. Maximum projections of confocal z-stacks are shown. (B) Flow chart of the heat shock and live-hemocyte imaging procedure. SPT starts 10 min after heat shock and continues over multiple cells (1-2 min per cell) for a total of 30 min with each sample. (C) Global chromatin-bound fractions for HSF-Halo and Halo-GAF at room temperature (*RT*) and *37.5°C*. Fast and slow-tracking results with propagated errors. ***, *p* < 0.001, unpaired *t*-test for fast-tracking (n=3-4). (D) Corrected average residence times for HSF-Halo and Halo-GAF at *RT* and 37.5°C. Error bars represent bootstrapped SD. (E) Diffusion coefficients derived using Spot-On for bound fractions of HSF-Halo and Halo-GAF at *RT* and *37.5°C*, and Halo-H2B at *RT. ***, p* < 0.001, unpaired *t*-test for fast-tracking (n=3-4). (F) Average MSD (mean ± SE) versus lag time of bound trajectories classified by vbSPT for HSF-Halo, Halo-GAF and Halo-H2B at *RT* and *37.5°C*, and Halo-H2B at RT. See Fig. S10A for a zoomed-in section for GAF and H2B.

We performed live-cell SPT on HSF-Halo in fast- and slow-tracking modes in the *P{PZ}Hsf^03091^/Hsf^3^* genetic background, using hemocytes cultured at RT or heat shocked at 37.5 °C **(Fig. 4B)**. As expected, the overall binding F_bound_ of HSF from fast-tracking increases substantially from 24.9% to 44.3% upon heat shock **(Fig. 4C** and **Fig. S8B)**. Importantly, two-component exponential decay fitting of the HSF-Halo survival curves derived from slow tracking reveals a substantial increase of stable binding F_sb_ from 5.9% to 14.2% upon heat shock with no measurable change of residence time τ_sb_ (47s) **(Fig. 4C-D** and **Fig. S8C, E)**. Thus, heat shock elevates the stable chromatin-binding HSF trimer fraction without affecting the dissociation rate (inverse of residence time), and suggests that the limited stable binding at RT (F_sb_ = 5.9%) is due to low-level trimerization^57^. Distinct from HSF dynamics, GAF shows a small overall reduction in F_bound_ on heat shock (from 77.3% to 69.6%), and similarly for the stable binding fraction F_sb_ (from 29.4% to 22.1%) **(Fig. 4C)**. Importantly, the residence time τ_sb_ for GAF remains unchanged after heat shock **(Fig. 4D and S8D-E)**.

### Chromatin-bound GAF displays H2B-like confinement

The diffusion coefficient of chromatin-bound HSF measured by fast-tracking (D_bound_, average of both stable- and transient-binding) at RT where HSF monomers predominate is ^~^4-fold greater than that of HSF trimers induced by heat shock (D_bound_= 0.075 vs 0.019 um^2^/s) **(Fig. 4E)**. HSF monomers also exhibit >10-fold larger D_bound_ values than GAF (D_bound_ = 0.0046 um^2^/s) **(Fig. 4E)**, indicating that a single DBD is more diffusive on chromatin than multiple DBDs. Intriguingly, D_bound_ for GAF approaches the H2B value (D_bound_ = 0.0020 um^2^/s) **(Fig. 4E)**. To strengthen these findings, we analyzed particle trajectories with vbSPT, a variational Bayesian Hidden Markov Model (HMM) algorithm that assigns bound and free diffusive states to individual particle displacements of each trajectory^63,64^ **(Fig. S9A-B)**. We classified particle trajectories as either ‘bound’ or ‘free’, excluding a small fraction showing two-state diffusivity **(Fig. S9A-D)**. The time-averaged MSD plots of the bound particles confirm that bound molecules move in small confined regions **(Fig. 4F and Fig. S10A)**, while free molecules undergo Brownian motion **(Fig. S10B)**. The MSD plot of chromatin-bound GAF trajectories reaches a plateau at low values, resembling that of H2B **(Fig. S10A)**, while HSF plateaus at higher values **(Fig. 4F)**. The radius of confinement (R_C_) gives median values for HSF monomers (0.13 μm), HSF trimers (0.10 μm), GAF multimers (0.07 μm), and H2B (0.06 μm) **(Fig. S10C)**. Together, the results indicate that chromatin-bound GAF is nearly as constrained as nucleosomal histones. Activated HSF trimers are less constrained than GAF, possibly due to fewer DBDs per complex and/or higher local chromatin mobility, but bound HSF monomers are more diffusive, consistent with the presence of only a single DBD.

### Constitutively high temporal occupancy defines pioneering activity

The steady-state open chromatin landscape at promoters and enhancers featuring nucleosome-depleted regions genome-wide^45,65,66^ belies highly dynamic interactions with transcription and chromatin factors^67–72^. The establishment and maintenance of chromatin accessibility, i.e. the sustained opening of chromatin, requires the joint activities of sequence-specific DNA-binding factors and ATP-dependent remodeling enzymes^12,19,24,25,73,74^, the latter proteins interacting with chromatin with a lifetime of seconds^67,69,70^. GAF directs pioneering functions not only by virtue of its affinity for nucleosomal DNA targets^12,75^ but also recruitment of chromatin remodelers^12,14,15,24,42,76^. Given the highly transient association and variable occupancy levels displayed by remodelers^67,70^, we hypothesized that GAF should instead sustain high occupancy along with protracted dwell time to continuously maintain open chromatin at cognate targets.

Temporal occupancy, the percent time of any duration for which a cognate site is factor-bound, depends on the number of GAF molecules (N_molecules_) per cell and the number of target sites (N_sites_) in the genome. In the context of the facilitated diffusion model^77^ in which transcription factors experience 3D nucleoplasmic diffusion, nonspecific binding, 1D diffusion, dissociation and re-binding until site-specific chromatin engagement, temporal occupancy is also dependent on the kinetics of target search and dissociation. Integration of our kinetic data from SPT with published genomic data allows calculation of temporal occupancy for GAF.

ChIP-seq identifies 3622 high-confidence GAF peaks from the hemocyte-like S2 cell line^17,78,79^. Similar numbers of GAF peaks are found by ChIP-seq analysis of larval imaginal tissues and embryos although the peaks from different cell types overlap partially ^25,80^. We measured GAF abundance by fluorescence flow cytometry using a calibrated CTCF standard^81^, estimating N_molecules_ = 56,683 ± 6,025 GAF molecules per hemocyte **(Fig. S11A)**. Given that circulating hemocytes are largely in the G2 cell cycle phase **(Fig. S11B-C)**, we estimate N_sites_ = 3622 × 4 genome copies = 14,488. From the overall chromatin-binding fraction [F_sb_], residence times and fractions for stable and transient chromatin-binding (τ_sb_, f_sb_, τ_tb_, f_tb_), we derived the average search time (τ_search_= 150 s, the time from GAF dissociation from one stable target to association with the next) and the sampling interval (SI = 71.5 s, the time from the start of one stable-binding event to the next stable event on the same chromatin target; see methods), assuming that GAF binds stably at specific sites and transiently elsewhere **(Fig 5A-B)**.

**Figure 5.**
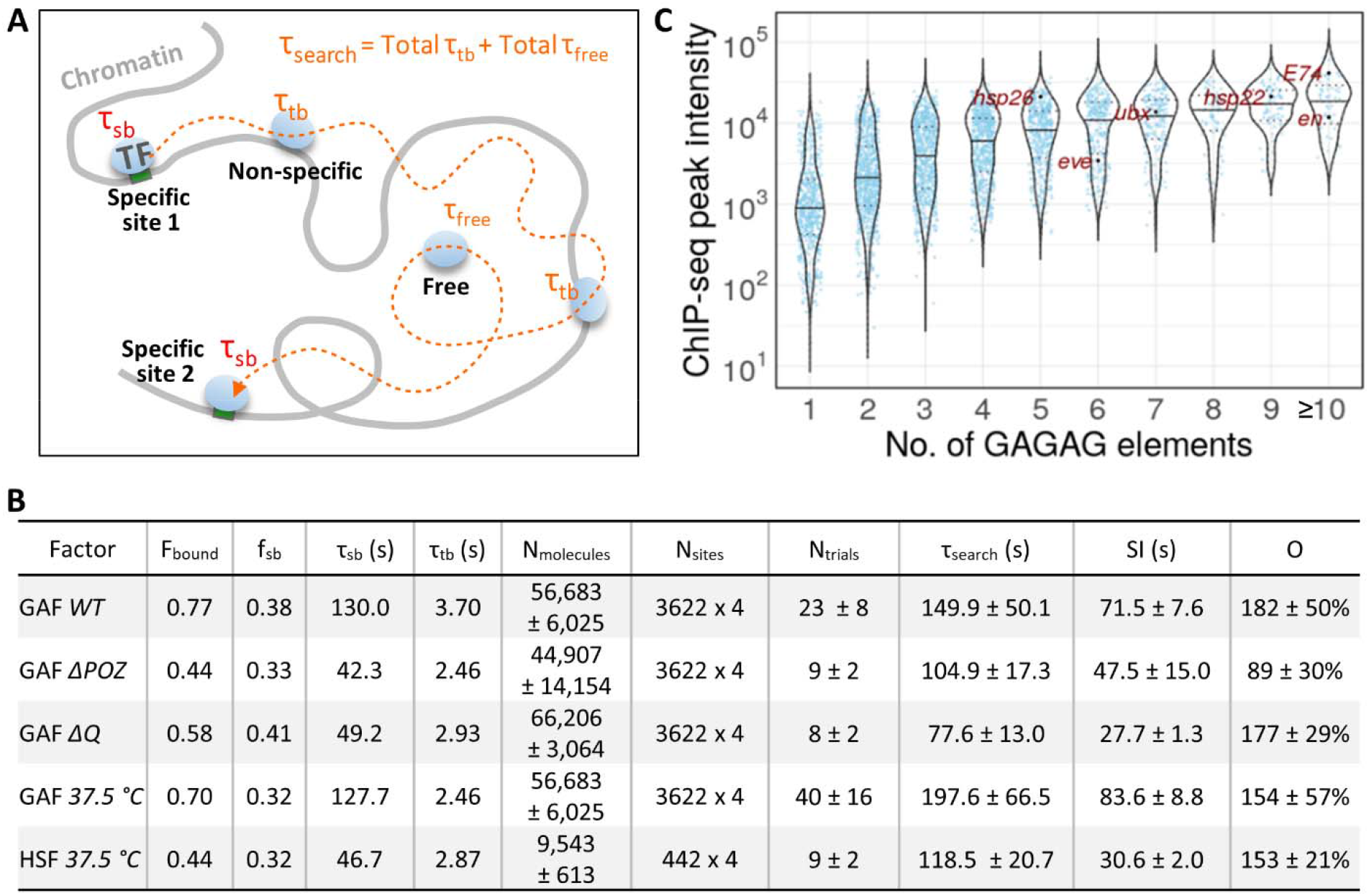
High site occupancy and remodeler autonomy quantifies pioneering criteria. (A) Schematic of a TF trajectory between two specific chromatin targets showing the search time τ_search_, stable τ_sb_ and transient τ_tb_ dwell times, and τ_free_. The TF molecule dissociates from a specific target, and samples nonspecific sites for a number of trials before encountering the next specific target. The τsearch equation is indicated (see methods). (B) Violin plots of GAF ChIP-seq peak intensities (analysis of S3 Table from Fuda et al, 2015) in hemocyte-like S2 cells plotted by the number of non-overlapping GAGAG elements identified with HOMER^96^. (C) Key SPT and N_molecules_ parameters measured in this study and N_sites_ from the literature^17,48^ are used to calculate occupancy levels for GAF and HSF.

We calculated the average occupancy (occupancy = τ_sb_/SI) at 182% for a GAF target under nonshock conditions and at 154% after heat shock, assuming no change in the number of GAF molecules and targets. The occupancy for *ΔQ* (N_molecules_ = 66,206 ± 3,064) remains similar at 177% and is reduced to 89% for *ΔPOZ* (N_molecules_ = 44,907 ± 14,154) **(Fig. 5B)**. The 182% occupancy for *WT* GAF is averaged over the 3622 GAF peaks whose intensities are correlated with increasing numbers of GAGAG elements. Notably, ^~^65% of GAF peaks harbor more than two non-overlapping GAGAG elements, with median peak intensity rising to a plateau at 6-7 clustered elements^17^(Fig. 5C), e.g. at *ubx, engrailed, E74, eve*, and *Hsp* genes^4^. The results indicate that GAF binds at least as a dimer on average, with a distribution tending towards larger oligomers for peaks showing high ChIP-seq signals, which can be attributed to the cooperative binding of GAF, as demonstrated in vitro^39,40^. At this subset of highly-enriched sites, GAF may bind as a multimeric complex with essentially full temporal occupancy despite factor on-off dynamics **(Fig. 6)** for a time period of any duration in which GAF levels and the number of GAF targets remain unchanged.

**Figure 6.**
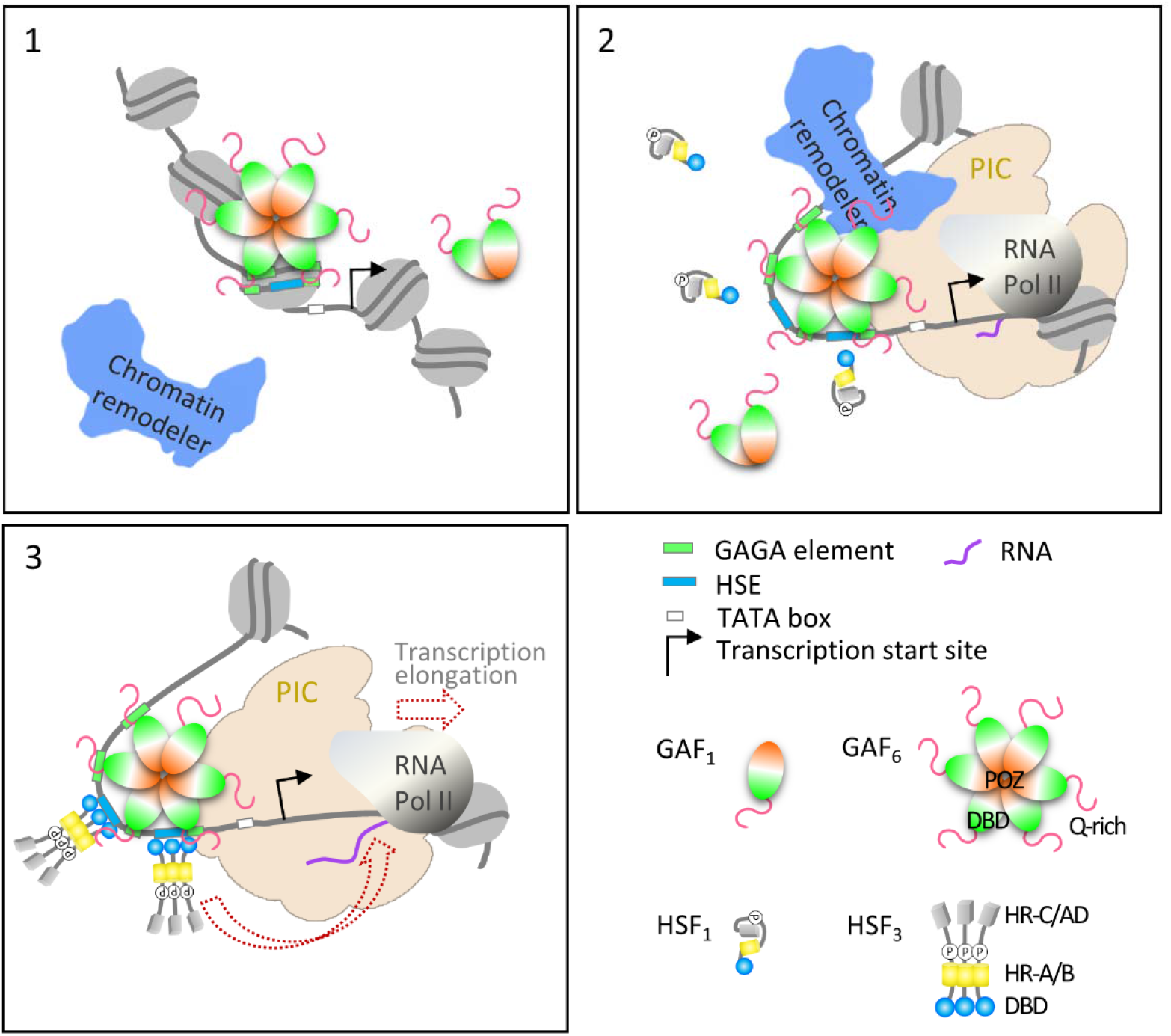
Pioneering of chromatin accessibility is a process involving multiple inputs. Model: GAF binds autonomously to nucleosomal sites at the first stage of pioneering (box 1). High GAF occupancy at a *Hsp* promoter with clustered GAGA elements maintains chromatin accessibility for neighboring factor HSF and assembly of the preinitiation complex and paused RNA Pol II (box 2). The substantially constrained diffusivity (D_bound_) of stably bound GAF may reflect multisite interactions of a GAF multimer with clustered GAGA elements (Katsani et al, 1999), locking down GAF with prolonged residence time. HSF trimers bind to accessible chromatin DNA with high affinity on heat shock to trigger RNA Pol II elongation (box 3). See discussion for details.

We note that many GAF-binding peaks overlap with Pipsqueak, a related POZ-domain transcription factor^82,83^ and partially with CLAMP, another GA repeat-binding pioneer factor in *Drosophila*^84,85^. The overall site occupancy at GAF locations on chromatin is therefore likely to be further supplemented when the contributions of Pipsqueak and CLAMP are quantified.

For HSF, we determined N_molecules_ = 9,543 ± 613 for a sole source, transgenic HSF-Halo under natural expression control in the *P{PZ}Hsf^03091^/Hsf^3^* background **(Fig. S11A)**. A similar calculation for HSF-Halo binding to 442 genomic sites after heat shock^48^ gives τ_search_ = 119 s, and SI = 31 s, which results in an average HSF occupancy of 153%, or 51% for HSF trimers as the predominant species induced after 10-40 min heat shock **(Fig 5B)**. At highly-enriched HSF locations such as the major *Hsp* genes harboring several HSEs, it follows that one or more HSF trimers may occupy the promoter 100% of the time on full induction to release the paused RNA Pol II and rapidly recruit additional enzymes for a burst of transcription until system attenuation (**Fig. 6)**.

## Discussion

### Full complement of GAF domains promotes kinetics of stable chromatin association

Our single-particle tracking of GAF in live hemocytes reveals that GAF binds chromatin with an exceptionally long residence time τ_sb_ of ^~^2 min compared to other factors. Systematic mutagenesis showing that the DBD, the POZ multimerization domain and the Q-rich domains are required for viability also found that not just the DNA-binding Zinc finger, but all three GAF domains are required for its stable chromatin association and long τ_sb_ **(Fig. 2B-C)**. Of interest, the Zinc finger mutants *ZF^9^* and *ZF^10^* have a substantial residual slow-diffusing fraction in hemocytes, but show no specific binding on polytene chromosome bands **(Fig. 2B, D)**. This is possibly due to non-specific association of the altered zinc finger and the remaining basic regions of the DBD **(Fig. 2A and Fig. S4)**, and/or protein-protein interactions of the intact POZ or Q-rich IDR domains (<u>i</u>ntrinsically <u>d</u>isordered <u>r</u>egions). Overall, our findings are consistent with the contributions to specific and non-specific DNA binding shown by mutant DBDs of mammalian TFs ^32,36,86^, and with the observation of IDR-assisted, in vivo DNA binding specificity for two yeast TFs ^87^. Of note, mutant GAF proteins *ZF^9^* and *ZF^10^* also exhibit an aberrant, slow-diffusing species whose underlying mechanism is unknown and remains to be further explored **(Fig. S5F)**.

The POZ domain is found at the N-terminus of vertebrate and invertebrate transcriptional regulators implicated in development and disease^88^. Functionally, the POZ domain is involved in protein homo- and hetero-dimerization, as well as multimerization^88^. The POZ domain mediates multimerization of GAF, which facilitates cooperative binding to closely clustered GAGAG elements^39,40^, and assists long-distance promoter-enhancer interactions between well-separated GAGAG clusters^89^. As judged by the kinetic behaviors of the GAF POZ mutant, we conclude that multimerization of GAF constitutes a critical element for its ability to pioneer open chromatin. Similarly, a variant glucocorticoid receptor (GR) that mimics allosterically induced GR tetramerization converts GR to a super-receptor that enhances chromatin occupancy at normally inaccessible sites^90^.

### Autonomy from recruited chromatin remodelers

The coupling of GAF-mediated pioneering of chromatin accessibility to ATP-dependent chromatin remodeling activities has been reported from the outset of studies on the mechanism underlying DNase hypersensitive sites. Biochemical experiments have since demonstrated that GAF directly recruits remodelers NURF and PBAP via protein-protein interactions, in addition to a number of other chromatin-based factors^76^. Using mutants for NURF and PBAP, we now show that their recruitment is not obligatory for GAF to kinetically engage chromatin targets. This indicates that GAF is largely autonomous from recruited remodelers and that the ensuing chromatin remodeling to antagonize competing processes of nucleosome encroachment primarily benefits the binding or activity of neighboring TFs to chromatin targets. While other factors or the global background of remodeling activities are not excluded from modulating GAF binding, GAF’s relative autonomy from two prominent recruited members of the SWI/SNF and ISWI remodeler families at the initial step of chromatin association may define an important property of transcription factors that act as pioneering agents.

### High constitutive and inducible temporal occupancies by GAF and HSF

By curating genomic databases for the number of genomic targets and integrating these parameters with measured kinetics and abundance, we found that GAF binds to its target sites with temporal occupancy of 182% for 3622 high-confidence ChIP-seq peaks^17^ **(Fig. 5A-C)**, i.e. with near full occupancy as a dimer on average. For genomic sites with greater than average GAF ChIP enrichment and number of GAGAG elements **(Fig. 5B)**, this occupancy is likely to involve higher oligomers, consistent with the native biochemical states of GAF complexes. Such high occupancy, or kinetic persistence, whereby factor dissociation from chromatin and replacement are essentially simultaneous, maintains a constant barrier and magnet for remodeler recruitment at GAF targets. (GAF multimers appear as stable biochemical complexes^39,40^, but dynamic exchange of GAF monomers within a multimeric complex is possible in principle and awaits further study). We envision that a substantial fraction of GAF targets in the genome displays high oligomeric status and temporal occupancy (^~^100%), while the remaining targets showing progressively lower occupancies, consistent with the genome-wide continua of transcription factor binding levels on metazoan genomes that reflect functional, quasi-functional, and nonfunctional transcription control^91,92^.

Unlike GAF, HSF monomers under room temperature conditions inducibly trimerize on heat shock^57^ which significantly increases the chromatin-bound fraction (F_bound_) without changing stable residence time (τ_sb_). For ^~^400 activated HSF-binding sites on the *Drosophila* genome^48^, we estimate an average of ^~^50% temporal occupancy by one HSF trimer. At the major *Hsp* loci harboring greater than average ChIP-seq signal intensity and multiple HSE elements, we envision that one or more HSF trimers engage at near full (100%) temporal occupancy. Full occupancy by HSF on heat shock induction may be required to facilitate release of the paused RNA Polymerase II previously established by GAF, and to sustain recruitment-release of new transcription pre-initiation complexes for a strong transcriptional burst of HS-responsive genes.

### Pioneering of chromatin accessibility is a process involving multiple inputs

The pioneer transcription factor concept has been introduced and elaborated for over two decades with a focus on special nucleosome binding properties of the FoxA1 prototype proposed to initiate establishment of chromatin accessibility for the benefit of consequent binding of neighboring TFs^93^. However, there continues to be debate whether the reported properties of FoxA1 are sufficiently distinct to set it apart from other transcription factors^16,35,94,95^. Our early findings on GAF that predate the controversy have documented that nucleosome binding and ATP-dependent remodeling are functionally coupled in a biochemical assay^12^, and additional studies to the present - including the genome-wide effects of remodeler depletion on chromatin accessibility^24^ and nucleosome positioning at GAF targets^73^ - support the concept that GAF directly recruits remodelers for the site-specific creation of accessible chromatin.

Thus, there is ample evidence to include remodeler recruitment by GAF or other TFs^35^ as a fundamental biochemical criterion for pioneering besides affinity for nucleosomes or closed chromatin^36^. Our finding of autonomous remodeler recruitment (chromatin interaction kinetics of GAF being largely unaffected in NURF and PBAP mutants) provides additional insight on the hierarchical nature of pioneering wherein GAF binding to chromatin at the initial stage of pioneering (stage 1) would be followed by remodeler recruitment and ATP-dependent nucleosome mobilization to create DNase hypersensitivity, thereby facilitating assembly of the transcription preinitiation complex (PIC), paused Pol II, and the inducible binding of HSF (stage 2) **(Fig. 6)**. In addition to remodeler recruitment, constitutively full temporal occupancy by GAF revealed by the single-particle kinetics provides a quantitative criterion for pioneering long-term chromatin accessibility primed for the transcriptional responses to homeostatic, environmental, and developmental signals.

However, we emphasize that high temporal occupancy is not an obligatory consequence of GAF’s long residence time on chromatin, or its multimeric, cooperative binding to GAGAG elements. Occupancy is also dependent on cellular GAF expression, abundance, and the number and genomic distribution of GAGAG elements in *Drosophila*. Thus, it may be instructive to consider pioneering as an active process with the multiple inputs we have described - autonomous factor-binding to closed chromatin, remodeler recruitment, nucleosome mobilization, and for the subset of transcription factors that maintain accessible chromatin constitutively, correspondingly high temporal occupancy, not excluding additional criteria to be identified. We hope that inclusion of these biochemical and kinetic principles guides further investigations on the substantial fraction of computationally identified human TFs (16% of ^~^700 TFs)^16^ that may pioneer chromatin accessibility as a basic mechanism of genome regulation in eukaryotic organisms.

## Methods

### Fly strain construction

#### CRISPR/Cas9-mediated genome editing

HaloTag was inserted downstream of the start codon of endogenous *Trl* via homology-directed repair (HDR) and CRISPR/Cas9 to generate the Halo-GAF knock-in strain **(Fig. S1 A-C)**. The donor repair template was constructed on the pScarlessHD-DsRed plasmid (a gift from Kate O’Connor-Giles, Addgene plasmid # 64703), which contains a DsRed selection marker cassette flanked by PBac transposon ends and TTAA sites^97^. The donor plasmid was designed such that after HDR, the DsRed cassette is inserted into a nearby genomic TTAA site adjacent to the gRNA target in the coding region that is close to the ATG start codon. Approximately 1 kb downstream of the gRNA site was cloned as the right homology arm (RHA), with silent mutations introduced to destroy the gRNA sequence in the donor plasmid. Similarly, 1kb upstream of the genomic TTAA site was cloned as the left homology arm (LHA). LHA and RHA mediate HDR upon Cas9 cleavage, inserting HaloTag along with the DsRed cassette. Flies that underwent HDR were identified with DsRed eye fluorescence. The DsRed cassette was removed with a single cross to a fly strain expressing PBac transposase, as indicated by loss of fluorescence, leaving only one TTAA site, thus allowing scarless HaloTag knock-in with a removable selection marker. A flexible linker GGSGS was added between Halo and GAF. The HaloTag knock-in was verified by fluorescent staining and DNA sequencing.

After constructing the Halo-GAF strain, deletions of the Halo-GAF fusion protein were generated by CRISPR-Cas9 gene editing. A 90 bp (30 AA, Δ90-119) precise deletion was generated in the POZ domain (*ΔPOZ*) by HDR. Small deletions in the zinc finger of the DBD (*ZF^9^*, R356Δ, N357Δ; and *ZF^10^*, R356Δ) were generated by random indels. For ΔQ, two gRNAs targeting the Q-rich domains of the long or short GAF isoforms were introduced at the same time to screen for indels creating frameshifts and truncations of both Q-rich domains. To screen for desired mutants, lethal or reduced viability strains were selected and characterized by PCR and DNAsequencing.

All gRNAs were cloned into pCFD5. Donor and gRNA plasmids were mixed to a final concentration of 200ng/uL and 500^~^600ng/uL, respectively, and injected into fly strains expressing Cas9 in the germline (yw;nos-Cas9(II-attP40), a gift from NIG-FLY, Japan). To generate Halo-GAF mutants, the Halo-GAF knock-in strain was crossed to yw;nos-Cas9(II-attP40) for injection. All fly embryo injections were performed by BestGene Inc (CA).

#### Transgenic fly construction via PhiC31 integrase

Trl gene and ^~^1kb flanking genomic sequence was cloned from a BAC genomic clone (BACR11B23) into pattB (backbone taken from pattB-aubergine-ADH-gf, Addgene plasmid # 69448, a gift from Phillip Zamore) via recombineering^98^. HaloTag was inserted upstream of the stop codon for the long and short isoforms, respectively, along with a removable Cam^R^ selection cassette via recombineering^99^. The CamR cassette was flanked by the 8-bp NotI restriction sites (plus an additional bp to ensure that HaloTag is in frame) and removed by NotI digestion and re-ligation, leaving a GGSGSAAA linker sequence between GAF and HaloTag. The constructs were incorporated into the attP2 site in the *Drosophila* genome via PhiC31 integrase^100^ (Fig. S1E), generating the GAF^L^-Halo and GAF^S^-Halo transgenic strains, which express a Halo-tagged long or short GAF isoform and the other untagged isoform. The functionality of recombinant fusion proteins was verified by rescue of the lethal alleles *Trl^13C^/Trl^R67 3^*. We similarly generated the HSF-Halo transgenic fly strain at the attP2 site from the genomic clone BACR33K09. The functionality of HSF-Halo was verified by rescue of *P{PZ}Hsf^03091^/Hsf^3^* lethal alleles^58,59^. The Halo-H2B transgenic strain was similarly constructed at the attP2 site, with a ^~^4.9 kb DNA fragment containing five *Drosophila* histone genes for HaloTag insertion at the N-terminus of H2B.

#### Mutant fly strains

Mutant alleles were obtained from the Bloomington *Drosophila* Stock center (BDSD, IN): *Trl^13C^*(BDSC:58473); *Trl^R67^* (BDSC:58475); *P{PZ}Hsf^03091^* (BDSC:11271); *Hsf^3^* (BDSC:5488); *P{EP}Bap170^G5986^* (BDSC:28471); *Bap170^Δ135^* (BDSC:63807); *Nurf301^4^* (BDSC:9904); and *Df(3L)Exel6084* (BDSC:7563).

Genotypes of fly strains in Fig. 3 and Fig. S6 (only GAF^L^-Halo isoform is shown, GAF^s^-Halo strains have the same corresponding genotypes except expressing GAF^s^-Halo):

*WT: p{GAF^L^-Halo}attP2*
*bap170: Bap170^Δ135^/P{EP}Bap170^G5986^; p{GAF^L^-Halo}attP2* (generated by crossing *Bap170^Δ135^; p{GAF^L^-Halo}attP2/T(2;3)TSTL, CyO: TM6B, Tb^1^* to *P{EP}Bap170^G5986^; p{GAF^L^-Halo}attP2/T(2;3)TSTL, CyO: TM6B, Tb^1^*)
*nurf301: Nurf301^4^, p{GAF^L^-Halo}attP2/Df(3L)Exel6084, p{GAF^L^-Halo}attP2* (generated by crossing *Nurf301^4^, p{GAF^L^-Halo}attP2/TM6B, Tb^1^* to *Df(3L)Exel6084, p{GAF^L^-Halo}attP2/TM6B, Tb^1^*)

Genotypes of fly strains in Fig. S8A:

*WT: p{HSF-Halo}attP2*
*trl: Trl^13C^, p{HSF-Halo}attP2/Trl^R67^, p{HSF-Halo}attP2* (generated by crossing *Trl^13C^, p{HSF-Halo}attP2/TM6B, Tb^1^* to *Trl^R67^, p{HSF-Halo}attP2/TM6B, Tb^1^*)
*bap170: Bap170^Δ135^/P{EP}Bap170^G5986^; p{HSF-Halo}attP2* (generated by crossing *Bap170^Δ135^; p{HSF-Halo}attP2/T(2;3)TSTL, CyO: TM6B, Tb^1^* to *P{EP}Bap170^G5986^; p{HSF-Halo}attP2/T(2;3)TSTL, CyO: TM6B, Tb^1^*)
*nurf301: Nurf301^4^, p{HSF-Halo}attP2/Df(3L)Exel6084, p{HSF-Halo}attP2* (generated by crossing *Nurf301^4^, p{HSF-Halo}attP2/TM6B, Tb^1^* to *Df(3L)Exel6084, p{HSF-Halo}attP2/TM6B, Tb^1^*)

### Single-particle imaging in live *Drosophila* hemocytes

#### Sample preparation

Single-molecule live cell imaging was performed with 3^rd^ instar larval hemocytes, representing mainly plasmatocytes (>90% of *Drosophila* hemocytes). Hemocytes were released from 5-10 thoroughly washed larvae into a sample chamber containing 1 mL filtered Schneider’s *Drosophila* medium (Gibco™ 21720024) including an EDTA-free protease inhibitor cocktail (Roche 4693159001). The sample chamber is an Attofluor™ Cell Chamber (Invitrogen, A7816) assembled with round coverglass (Electron Microscopy Sciences, 72290-12) cleaned by flaming. Cells were stained with 0.2^~^1.5 nM JF552/JFX554 for 30 min at room temperature, during which hemocytes adhered to the coverglass bottom of the imaging chamber. During incubation steps, sample chambers were covered with aluminum foil to minimize evaporation and block light. Cells were then briefly washed twice with Schneider’s media without protease inhibitor and imaged on a custom-built wide-field SPT fluorescence microscope ^101^.

#### Single-particle imaging

All single-particle imaging were carried out on an Axio Observer Z1 microscope (Zeiss, Germany) equipped with an Plan-Apochromat 150x/1.35 glycerin-immersion objective (ZEISS, Germany), and a C9100-13 EM-CCD camera (Hamamatsu Photonics, Japan) featuring 512×512 pixels with 16 μm pixel size. The pixel size of recorded images is 16 μm/150 = 107 nm. JF552/JFX554 were excited with a CL555-100 555 nm laser (CrystaLaser, Reno, NV) and the emission light was passed through a filter cube containing a 561 nm BrightLine single-edge beamsplitter, a 612/69 nm BrightLine single-band bandpass emission filter (Semrock, Rochester, NY), then through a 750 nm blocking edge BrightLine multiphoton short-pass emission filter and a 405/488/561/635 nm StopLine quad-notch filter (Semrock, Rochester, NY) before entering the camera. The EM-CCD camera was operated at ^~^-80°C (forced-air cooling) and 1200x EM gain. We used the ZEN (ZEISS, Germany) and HCImage (Hamamatsu Photonics, Japan) software to operate the microscope and camera, respectively. Imaging was carried out at room temperature except for heat shock experiments in which samples were imaged in a stage-top incubator with temperature control (H301-MINI chamber with UNO-T-H controller, Okolab, Italy)

In the fast-tracking regime, time-lapse movies with a 128×128 pixel field of view were acquired with high laser power (^~^1 kW/cm^2^) and 10 ms exposure time for 1.5-2 min. Initial laser excitation leads to simultaneous emissions of all labeled molecules, marking locations of individual nuclei in the field of view. Cells with relatively homogenous initial nuclear glow (interphase) were imaged. Emitting molecules quickly enter the “dark” state and stochastically reemit. Cells were stained with JFX554, with the concentration optimized (1-1.5 nM) to achieve sparse single-particles per nucleus per frame after 10^~^30 s of the initial glow, minimizing mis-tracking.

For slow tracking, we used low laser power (^~^36 W/cm^2^) and imaged a 256×256 pixel field of view with 500 ms exposure time for 2-5 min. We labeled cells with 50 nM of a far-red dye JF700 to block most Halo-tagged proteins and at the same time adjusted a low concentration of JF552 to visualize only 2-10 molecules/frame. This sparse labeling approach allows tracking of chromatin-bound molecules with minimal photobleaching.

### Single-particle data processing and statistical analysis

#### Image pre-processing

Raw time-lapse data were pre-processed in Fiji^102^ to convert to 16-bit TIFF format and extract a substack with sparse single particles (<5 particles/nucleus for fast tracking, and < 10 particles/nucleus for slow tracking). A maximum-intensity Z projection of each movie was generated to outline cell nuclei as the ROI. A corresponding binary mask was created to isolate nuclear trajectories for subsequent analysis.

#### Single-particle localizing and tracking

Single particles were localized and tracked using the open-source program DiaTrack v3.05^103^. Tracking was performed with a 6 pixel (^~^0.65 μm) maximum jump allowance between consecutive frames for fast tracking. All factors we imaged showed displacements within this cut-off as informed by the frequency histogram **(Fig. S2B, S5A, S6A, and S7B)**. Since slow tracking selectively imaged bound particles, the maximum jump allowance between consecutive frames was set as 3 pixels (^~^0.32 μm) to minimize misconnection. HSF-Halo showed displacement histograms with a smooth tail within this range, while Halo-GAF and Halo-H2B displacements were mostly within 0.2 μm (not shown). For analysis of slow-tracking experiments to measure residence times, we allowed gaps in trajectories to account for blinking or missed localizations, with 3-frame maximum blinking and a more stringent 2-pixel maximum jump.

#### Analyzing fast-tracking data

We used a custom R package Sojourner (https://github.com/sheng-liu/sojourner) to extract trajectories from MATLAB files generated by DiaTrack, which contain information on x, y coordinates and frame number (time) that were applied for computation of kinetic parameters. Trajectories found within the nucleus were isolated (‘masked’ trajectories) using Sojourner and binary masks generated during pre-processing. Average length of trajectories was 11-21 displacements (median = 5-8).

##### Spot-On

We used Spot-on to perform two-state kinetic modeling of displacements from all ‘masked’ trajectories^33^, to derive diffusion coefficients (D_bound_, D_free_) and the corresponding fractions (F_bound_, F_free_=1-F_bound_). The Spot-On python package was used with the following parameters: bin width = 0.01 μm, number of timepoints = 6, jumps to consider = 4, max jump = 1 μm, gapsAllowed = 0, and Z correction with dZ = 0.6 μm. Mean and SD of F_bound_, D_bound_ and D_free_ were calculated from 3-5 biological replicates.

##### vbSPT (variational Bayesian single-particle tracking) HMM

A matlab program running vbSPT HMM^63,64^ (https://gitlab.com/anders.sejr.hansen/anisotropy) was modified to assign each trajectory displacement into two states, ‘bound’ or ‘free’. Then each trajectory was sub-classified as ‘bound’ if all displacements are classified as bound state; ‘free’ if all displacements are classified as free state; a small fraction of trajectories containing two states were omitted from related analysis. The sub-classified ‘bound’ and ‘free’ trajectories from biological replicates of the same conditions were pooled together and used to calculate mean squared displacements (MSD).

To calculate the apparent radius of confinement (Rc) for individual trajectories, we fit each MSD curve with the following confined diffusion model^36,104^,

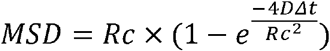

#### Analyzing slow-tracking data

Using the Sojourner R package, the apparent dwell times (temporal length of trajectories) were determined for all “masked” trajectories lasting at least 3 frames.

1-CDF curves were generated and fit to a double exponential decay model:

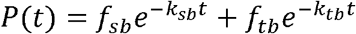

where k_sb_ and k_tb_ correspond to dissociation rates for stable- and transient-binding events, respectively; f_sb_ and f_tb_ correspond to the fraction of time the molecule spends at stable- and transient-binding sites, respectively, and f_sb_ + f_tb_ =1.

The survival curves reflect not only factor dissociation, but also photobleaching, axial and lateral cell or chromatin movements, fluctuating background, etc. To correct for all these factors, assuming that these processes affect Halo-H2B to the same extent as other proteins, and that bulk H2B dissociation is negligible in the experimental time frame of 2-5 min, we measured the apparent unbinding rate for Halo-H2B in the same way and used it as a correction factor for other proteins’ residence time^38^. The corrected average residence times for stable- (τ_sb_) and transient-binding (τ_tb_) were calculated as follows:

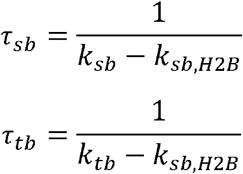

Mean and SD of k_sb_, k_tb_, f_sb_ and f_tb_ were calculated from 100 bootstrap samples, then mean and SD of k_sb_ and k_tb_ were used to calculate τ_sb_ and τ_tb_ with error propagation.

#### Calculating search kinetics and target occupancy

We integrated approaches from previous studies^32,105,106^ and calculated temporal occupancy as described^72^.

First, search time (τ_search_) is the average time it takes from a molecule dissociates off a specific site till it find the next specific site, during which the molecule samples nonspecific sites (each lasts for τ_tb_ on average) for a number of times (N_trials_) before encountering the next specific target (bound for τ_sb_ on average). τ_free_ is the average free time between two non-specific binding events. Assuming the molecule samples all non-specific and specific sites at random with equal accessibility, i.e., equal probability of binding to all sites, the average search time between two consecutive specific binding events is calculated as:

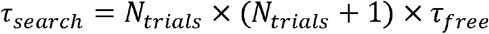

N_trials_ depends on the ratio of number of non-specific (N_ns_) to specific sites (N_s_), or *r_s_*:

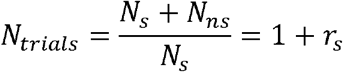

Thus,

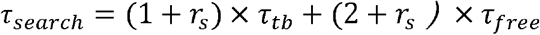

To determine r_s_, we considered two scenarios underlying detection of binding events during slow tracking^72^.

a. Blinking-limited (*r_s, bl_*): f_sb_ obtained by slow tracking is proportional to the fraction of time the molecule spends at specific sites (stable-binding) relative to the overall time it is bound to chromatin (stable- and transient-binding):

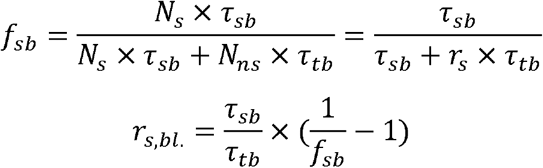
b. Diffusion-limited (*r_s, diff_*): assuming equal probability of binding to all sites, f_sb_ depends on the ratio of number of specific sites to all sites:

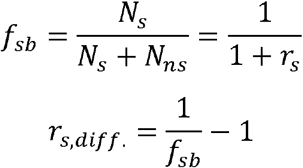

These physical processes can happen in the cell coincidentally and likely represent two extremes of a spectrum of behaviors single molecules can exhibit. We reasoned that the relative likelihood of detecting a blinking-limited binding event by slow tracking is proportional to the global fraction of bound molecules (F_bound_ obtained by fast tracking). Therefore, we computed a weighted average value for r_s_ as follows:

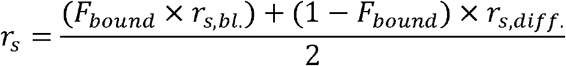

To obtain τ_free_, we considered that F_bound_ is proportional to the fraction of the time a molecule spends bound to chromatin either stably or transiently:

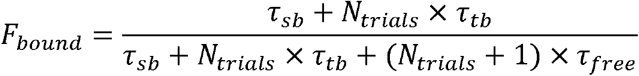

Thus,

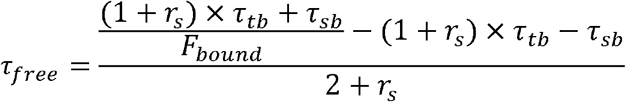

τ_search_ was calculated with the values derived for r_s_ and τ_free_, as shown above. We then estimated the sampling interval (SI), the average time between two consecutive binding events at a specific site^32^,

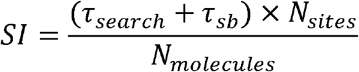

We used N_sites_ values presented by GAF and HSF ChIP-seq studies^17,48^. N_molecules_ was estimated by flow cytometry (see below). Finally, the average occupancy is the percentage of time a given specific site is occupied by the protein of interest:

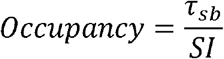

### Confocal microscopy

Hemocytes were prepared in the same way described above for live-cell imaging, except for staining with 50 nM JFX554 for 30 min at room temperature or heat shocked on a metal heat block at the indicated temperature (the time for heat shock overlaps with the final stage of staining so that dye labeling and heat shock were completed at the same time). During all incubation steps the sample chambers were covered with aluminum foil to minimize evaporation and block light. Then cells were briefly washed twice with PBS and immediately fixed in freshly made 4% formaldehyde (diluted from Pierce™ 16% Formaldehyde (w/v), Methanol-free, #28906) in 1X PBS for 15 min. Fixed samples were washed in PBS for 10 min, then stained with DAPI for 10 min and washed in PBS for 5 min. Sample chambers were filled with PBS and covered by a coverglass on top. Salivary glands were dissected, then stained and heat-shocked similarly as hemocytes except that JF554 staining was performed after fixation. Salivary gland samples were mounted on glass slides in VECTASHIELD® Antifade Mounting Medium (Vector Labs H-1000-10). We imaged these samples on a LSM800 Airyscan confocal microscope (Zeiss, Germany) with a 63X objective. Z-stacks were taken with 0.2 μm step size for hemocyte nuclei and 0.5 μm step size for salivary gland polytene nuclei. The same laser and scanning settings were used between samples in the same experiment.

### Estimation of cellular abundances for Halo-tagged proteins by flow cytometry

Cellular abundances of Halo-GAF and HSF-Halo were estimated by flow cytometry using a calibrated C32 CTCF-Halo U2OS human cell line^81,107^. U2OS cell samples were prepared as described^107^ with minor modifications. Briefly, we labeled WT and CTCF-Halo U2OS cells with 50 nM JF552 for 30 min at 37°C/5% CO_2_ in a tissue-culture incubator, washed out the dye (removed medium; rinsed with PBS, and incubated with fresh media for 5 min in the incubator) and then immediately prepared cells for flow cytometry. Resuspended cells were filtered through a 40 μm filter and placed on ice until their fluorescence was read out by the flow cytometer. To prepare fly hemocytes for flow cytometry, 20^~^30 3rd instar larvae were thoroughly washed and dissected on ice in a 1.5 mL eppendorf tube lid containing Schneider’s *Drosophila* medium (Gibco™ 21720024), EDTA-free protease inhibitor cocktail (Roche 4693159001) and 10% FBS (HyClone FBS SH30910.03, Cytiva, MA). Hemocytes were collected and stored on ice. After dissection was done for all fly strains, hemocytes were stained with 50nM JF552 in 1mL medium at room temperature for 30 min. All tubes were pre-rinsed with FBS to minimize cell stickiness to the tubes. After staining, hemocytes were centrifuged at 200 x g for 5 min, resuspended in 1 mL fresh medium and washed at room temperature for 15 min (longer wash time than for U2OS cells to remove non-specific cytoplasmic signals). The hemocytes were centrifuged again, resuspended in 500 μL medium and placed on ice until their fluorescence was read out by the flow cytometer.

We used a SH800 (Sony, Japan) cell sorter in analyze mode to measure fluorescence intensity in the U2OS cell lines and hemocytes with the same settings. Single live cells were gated using forward and side scattering. JF552 fluorescence was excited using a 561 nm laser and emission read out using a 617/30 band pass filter. The absolute abundance of protein of interest N_molecuels_ (mean number of molecules per cell) was obtained according to:

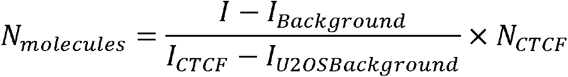

Where *I* is the average measured fluorescence intensity of the cells expressing the protein of interest, *I_Background_* is the average measured fluorescence intensity of hemocytes from a fly strain not expressing HaloTag (*w1118*), *I_CTCF_* is the average measured fluorescence intensity of the CTCF-Halo standard U2OS cell line, *I_U2OSBackground_* is the average measured fluorescence intensity of WT U2OS cells not expressing HaloTag, and *N_CTCF_* is the absolute abundance of CTCF-Halo (^~^109,800 proteins per cell). For each experiment, hemocytes and U2OS cells were stained with the same aliquot of JF552 stock solution and measured during the same flow cytometry session. We performed three to five biological replicates to get mean and standard deviation values.

### Characterization of cell cycle stage for larval hemocytes using the Fly-Fucci system

A fly strain with the Fly-Fucci markers under UAS control (UAS-CFP.E2f1.1-230 and UAS-Venus.NLS.CycB.1-266, BDSC:55122) was crossed to a hemocyte-specific driver Cg-Gal4 (BDSC:7011). 3^rd^ instar larvae were washed then dissected to release hemocytes in a drop of PBS on a slide, then the fluorescence of hemocytes were imaged immediately on a Zeiss Axioplan 2 compound microscope.

## ACKNOWLEDGEMENTS

We thank Anders Hansen and Maxime Woringer for discussion on Spot-On analyses, Anders Hansen, Claudia Cattoglio, Xavier Darzacq and Robert Tjian for the CTCF-Halo U2OS cell line and discussions on measuring factor abundance, Tim Lionnet, James Zhe Liu, Peng Dong, and Brian Mehl for advice on SPT, Yick Hin Ling and Sun Jay Yoo for assistance with data analysis, Pascal Vallotton for customization of DiaTrack software, Paul Badenhorst for advice on larval hemocyte culture, Steve Deluca for advice on bioinformatics, Weina Dai for imaging assistance, Erin Pryce and the Integrated Imaging Center for LSM 800 training, Yundong Liu for assembly and maintenance of a high performance computational platform, John Lis, Mike Levine, and Gordon Hager for discussion, and Wu Lab members for comments on the manuscript. This study was supported by HHMI funding to the TIC (C.W., Q.Z., and L.L.), Johns Hopkins Bloomberg Distinguished Professorship funds (C.W.), and National Institutes of Health grants GM132290-01 (C.W.) and DK127432 (C.W.).

## AUTHOR CONTRIBUTIONS

X.T. performed all genetic and imaging experiments with support from T.L., J.W., and Y.R. and all data analysis using R functions created by S.L. and X.T.. L.D.L. and Q.Z. synthesized JF552/JFX554 and JF700. X.T. and C.W. designed the study and wrote the paper with input from all authors.

## DECLARATION OF INTERESTS

L.D.L. and Q.Z. are listed as inventors on patents and patent applications whose values might be affected by publication.

## SUPPLEMENTARY FIGURES

**Figure S1.**
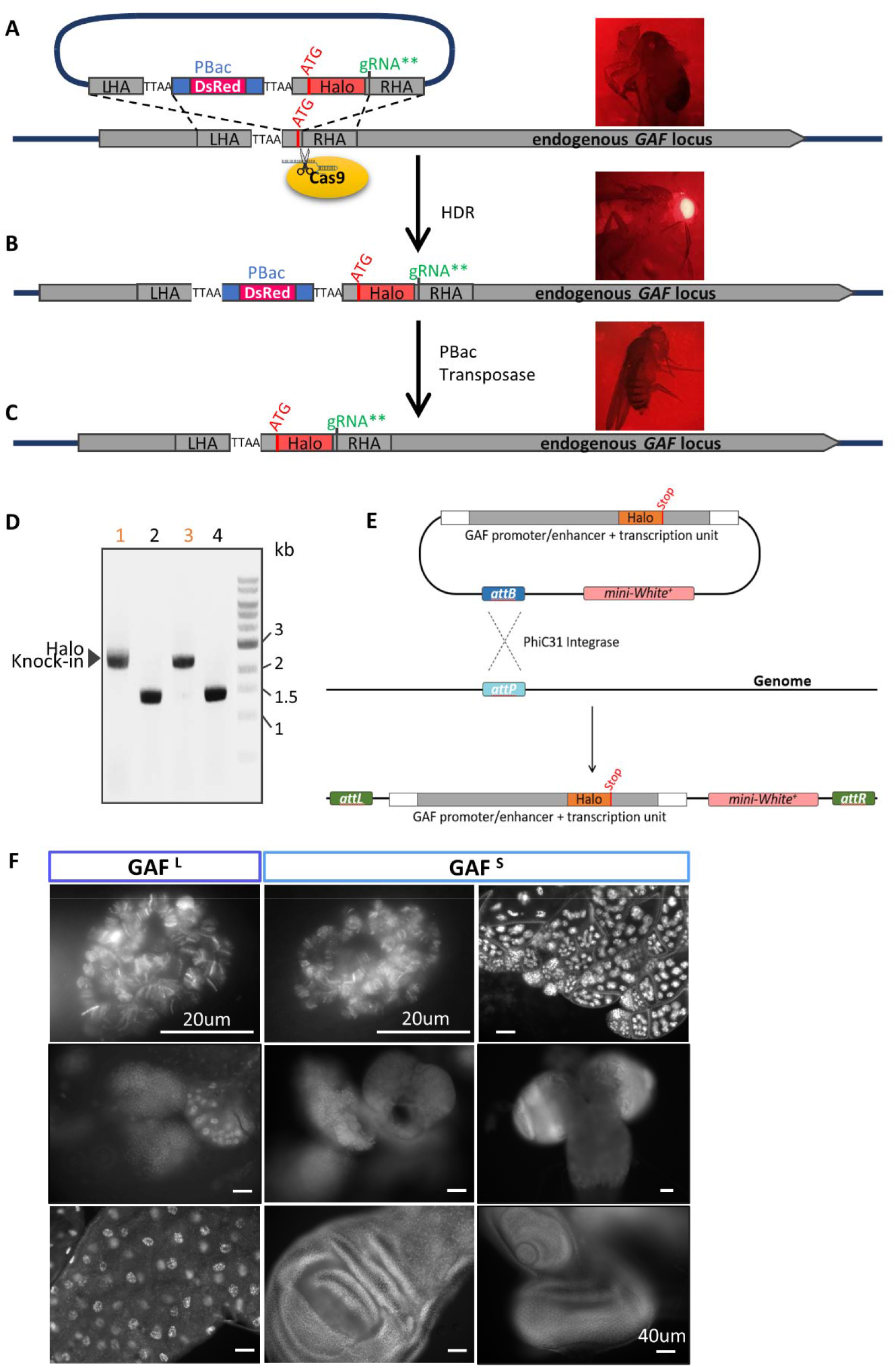
Generation of N-terminal Halo-GAF knock-in fly strain and C-terminal GAF-Halo transgenic fly strains. (A) Donor plasmid design for homology directed repair (HDR). HaloTag and a flexible linker (GGSGS, not shown) are placed downstream of the start codon ATG. A PBac transposon containing a DsRed cassette is inserted into a nearby genomic TTAA site adjacent to the gRNA target site in the coding region that is close to the start codon ATG. The TTAA site is duplicated so that both ends of the PBac transposon contain a TTAA sequence. Approximately 1 kb fragment downstream of the gRNA target site is cloned as the right homology arm (RHA), with silent mutations (gRNA**) introduced to destroy the gRNA PAM sequence in the donor plasmid. Similarly, a 1kb fragment upstream of the genomic TTAA site is cloned as the left homology arm (LHA). (B) LHA and RHA mediate HDR upon Cas9 cleavage, inserting HaloTag along with the DsRed cassette. Flies that have undergone HDR can be identified by eye DsRed fluorescence. (C) By crossing to a fly strain expressing PBac transposase, the DsRed cassette can be removed, as indicated by loss of fluorescence, leaving only one TTAA sequence, thereby allowing scarless HaloTag knock-in with a removable selection marker. Arrows indicate positions of the primers used for validating HaloTag insertion. (D) PCR validation of HaloTag knock-in after DsRed cassette removal (lane 1 and 3). Halo-GAF homozygous flies are viable, showing only 1 band ^~^900 bp larger than flies without HaloTag knock-in (lane 2 and 4). (E) Strategy used to generate transgenic fly strains GAF^L^-Halo, GAF^S^-Halo, Halo-H2B and HSF-Halo. An ^~^15 kb fragment containing the Trl transcription unit and ^~^1kb upstream and downstream regions was cloned, and HaloTag ORF was inserted upstream of the stop codons for GAF^L^ or GAF^S^, respectively. Thus, each of the two transgenic flies express a Halo-tagged GAFL or GAFS isoform and another isoform (non-tagged), under native Trl promoter control. (F) Tissue-specific expression of transgenic GAF^L^-Halo and GAF^S^-Halo. Shown are major expressing larval tissues. GAF^L^-Halo: salivary gland, lymph gland, intestine; GAF^S^-Halo: salivary gland, lymph gland, wing disc, ovary, brain, eye-antenna disc.

**Figure S2.**
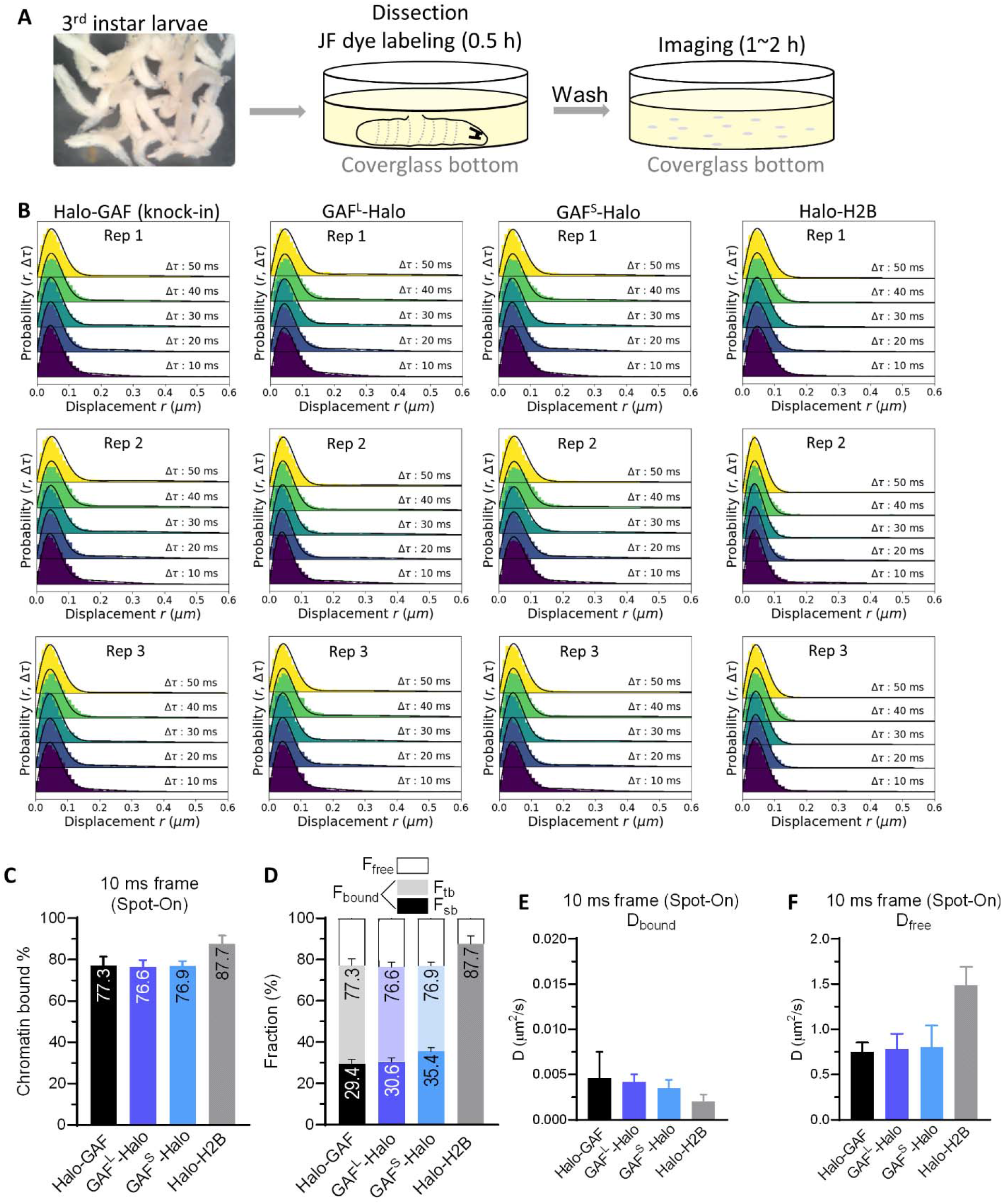
Hemocyte imaging and fast-tracking diffusive parameters for Halo-GAF, GAF^L^-Halo, GAF^S^-Halo, Halo-H2B. (A) Experimental timeline of single-particle imaging with 3rd instar larval hemocytes. 3rd instar larvae are washed with DI H2O (left) and dissected in a coverglass bottom dish containing Schneider’s medium and JF dye at room temperature. Upon dissection hemocytes are released into the medium and labeled for 30 min, the rest of the larval tissues are discarded (middle). Cells are briefly washed twice with fresh media and imaged for 1-2 h. (B) Spot-On fits of Halo-GAF, GAF^L^-Halo, GAF^S^-Halo, Halo-H2B fast-tracking data. (C) Spot-On kinetic modeling of fast-tracking data shows 77% of Halo-GAF is chromatin bound. Similar values are obtained for isoforms GAF^L^ and GAF^S^ individually tagged in the presence of untagged GAF isoforms. Results are mean ± SD from three biological replicates. (D) Chromatin-free fraction (F_free_), long- and short-lived chromatin-binding fractions (F_sb_ and F_tb_) of HaloTagged GAF fusions extracted from fast- and slow-tracking data in (C) and (Fig. S3E), respectively, with error propagation. (E) Diffusion coefficients of bound fraction (D_bound_) for Halo-GAF, GAF^L^-Halo, GAF^S^-Halo, Halo-H2B derived by Spot-On. (F) Diffusion coefficients of free fraction (Dfree) for Halo-GAF, GAF^L^-Halo, GAF^S^-Halo, Halo-H2B derived by Spot-On.

**Figure S3.**
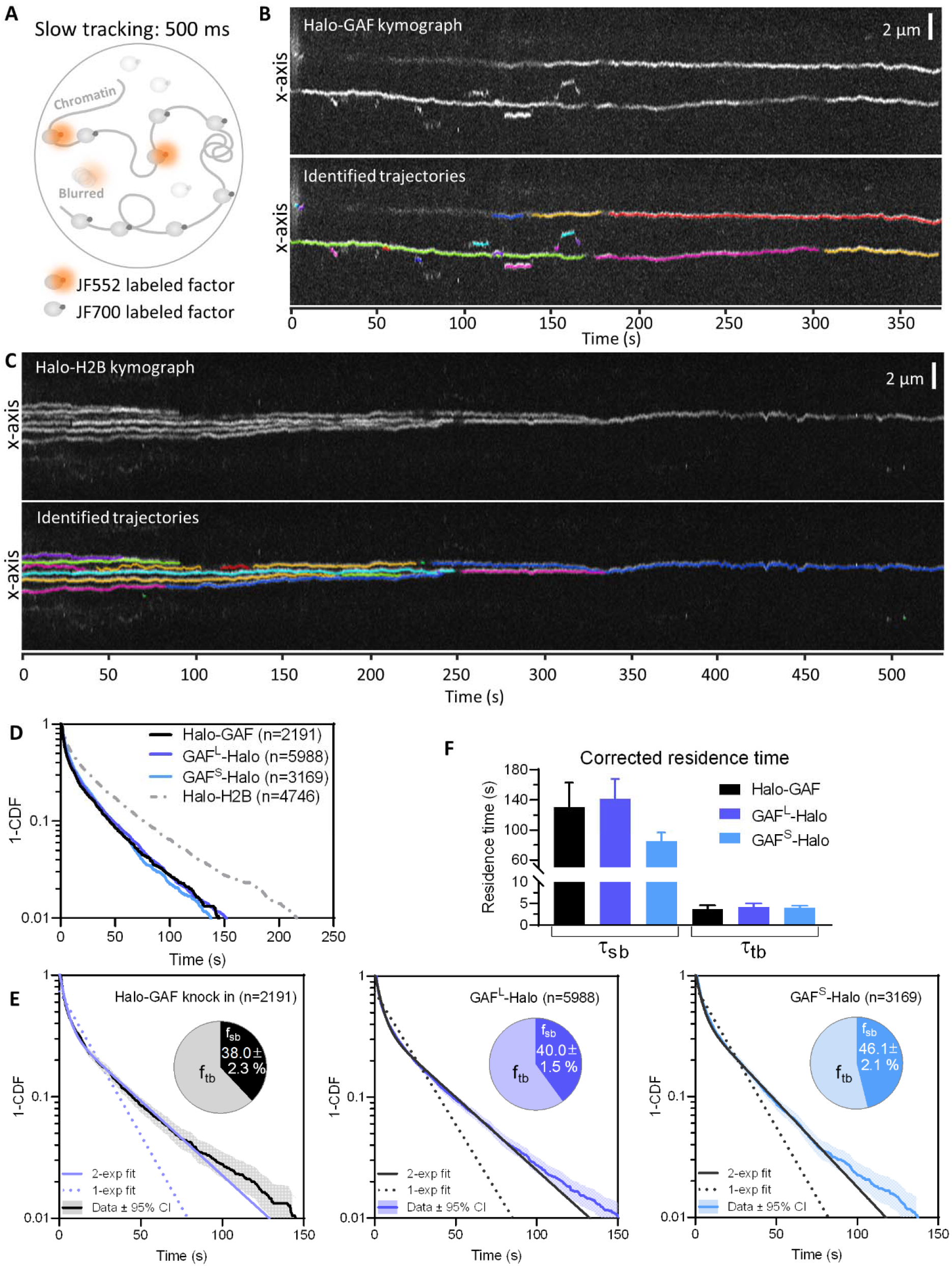
Slow tracking results for Halo-GAF, GAF^L^-Halo, GAF^S^-Halo, Halo-H2B. (A) Fast and slow tracking regimes. Fast tracking with 10 ms frame rate and high laser power allows single molecule imaging to distinguish slow (chromatin-bound) and fast (chromatin-free) diffusing subpopulations. Slow tracking uses low-intensity excitation and 500 ms exposure time to motion blur diffusing molecules and selectively observe the dwell times of chromatin-bound molecules. A higher concentration of JF700 is added to block labelling of most HaloTag protein fusions, while a much lower concentration of JF552 is used to sparsely label a small fraction of HaloTag so that each nucleus shows only 2^~^10 molecules per frame during image acquisition. (B) Kymograph of a Halo-GAF slow tracking movie shows traces of bound GAF molecules over time (upper). Trajectories identified from the raw movie are plotted on the kymograph using separate colors (lower). (C) Kymograph of a Halo-H2B slow tracking movie shows traces of bound H2B molecules over time (upper). Trajectories identified from the raw movie are plotted on the kymograph (lower). (D) Survival probability curves (1-CDF) plotted from apparent dwell times of thousands (n) of single-molecule chromatin-binding events for Halo-GAF, GAFL-Halo and GAFS-Halo. (E) One-component and two-component exponential fit of survival probabilities (1-CDF) from slow tracking data of Halo-GAF, GAF^L^-Halo and GAF^S^-Halo. Pie charts show the stable-binding (f_sb_) and transient-binding (f_sb_) fractions derived from two-component fits, and errors represent bootstrapped SD. (F) Corrected average residence times for stable- (τ_sb_) and transient- (τ_tb_) binding by transgenic GAF^L^-Halo and GAF^S^-Halo.

**Figure S4.**
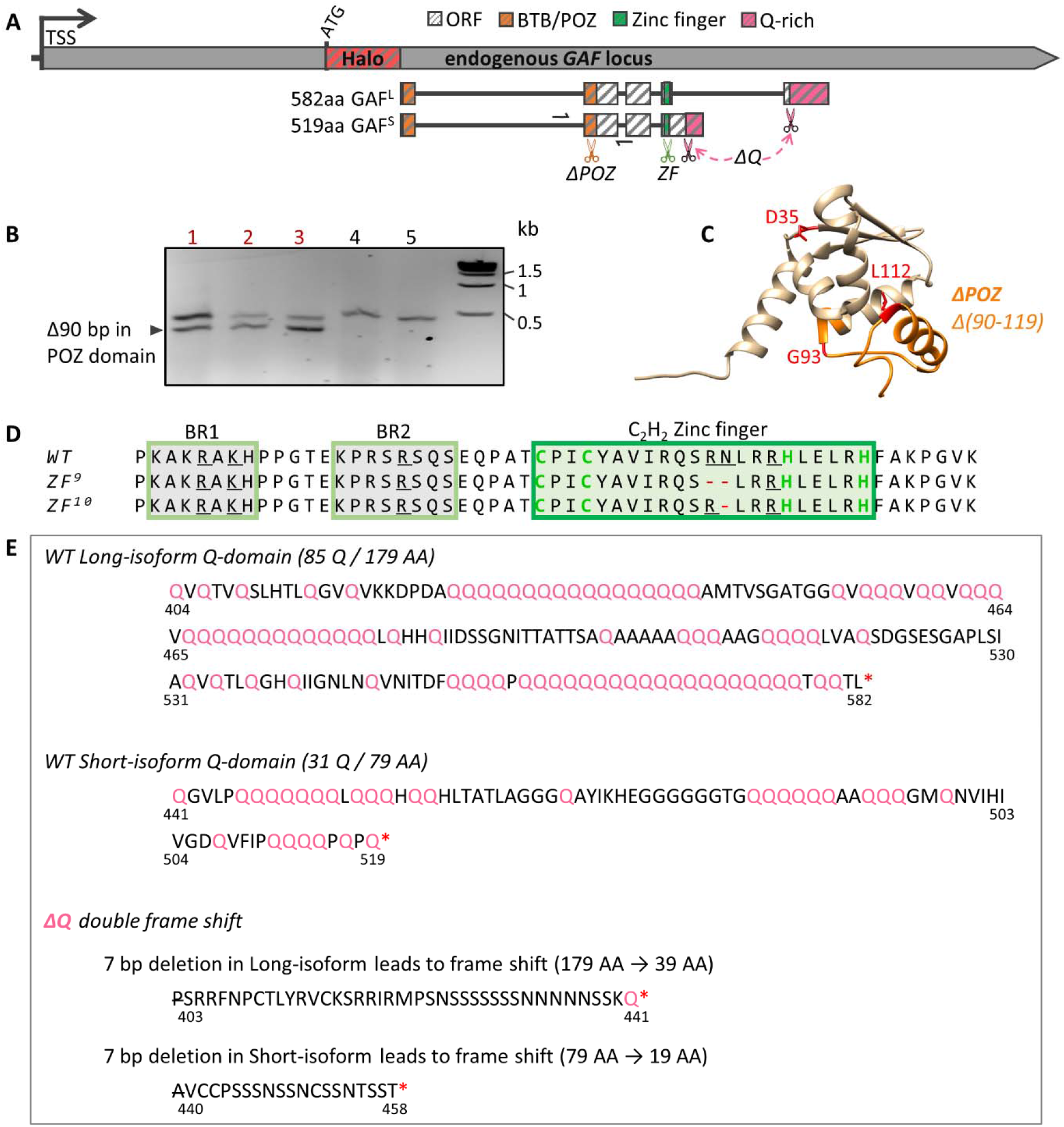
Generation of mutations in functional domains of Halo-GAF by CRISPR/Cas9 gene editing. (A) In the Halo-GAF fly strain, Cas9 and gRNA were introduced to target the BTB/POZ domain, zinc finger, and Q-rich domains, respectively. The BTB/POZ domain is separated by a large intron. A gRNA target site in the second exon (orange scissors) was selected and a donor plasmid containing a 90 bp deletion (*ΔPOZ*) was constructed for homology-directed repair (HDR). For zinc finger mutations, we selected a gRNA target site in the zinc finger coding region (green scissors), and screened for in-frame small deletions generated by non-homologous end joining (NHEJ). To generate deletions of both Q-rich domains in long and short isoforms (ΔQ), two gRNAs targeting the upstream ends of two Q-rich domains (pink scissors) were introduced at the same time, and we screened for double frame-shift deletions induced by NHEJ. After gRNA and donor plasmid injection, fly progeny were crossed with balancer chromosome, candidates homozygous-lethal or with reduced viabilities were verified by PCR and sanger sequencing. Half arrows indicate positions of the PCR primers used in (B). TSS, transcription start site. (B) PCR validation of ΔPOZ. Lanes 1-3 show two PCR bands indicating precise deletion in one allele; lanes 4-5 are two lines without the precise deletion. Sanger sequencing verified a precise 90 bp deletion in one allele. (C) POZ domain model structure. 90bp deletion in the second exon generates a 30-AA deletion (*Δ90-119*) of the POZ domain (ΔPOZ, orange), which includes G93 and L112 (red) that have been shown to be essential for transcription activation^29^. (D) Amino acid sequence of GAF DNA binding domain, which contains a single C2H2 zinc finger (green rectangle) and two upstream basic regions (BR1 and BR2, yellow rectangle). Amino acids involved in recognizing the GAGAG consensus sequence are underlined^108^. Two zinc finger mutations were isolated and verified by sanger sequencing, *ZF^9^* and *ZF^10^*, with R356 and N357, or R356 deleted, respectively. (E) Amino acid sequence of GAF Q-rich domains for long and short isoforms. In ΔQ, a 7bp deletion was identified by sanger sequencing in both isoforms at the upstream ends of Q-rich domains, resulting in frameshifts and truncations of the Q-rich domains from both isoforms. P403 in the long isoform and A440 in the short isoform are deleted and the subsequent amino acids are newly introduced by the frame shifts.

**Figure S5.**
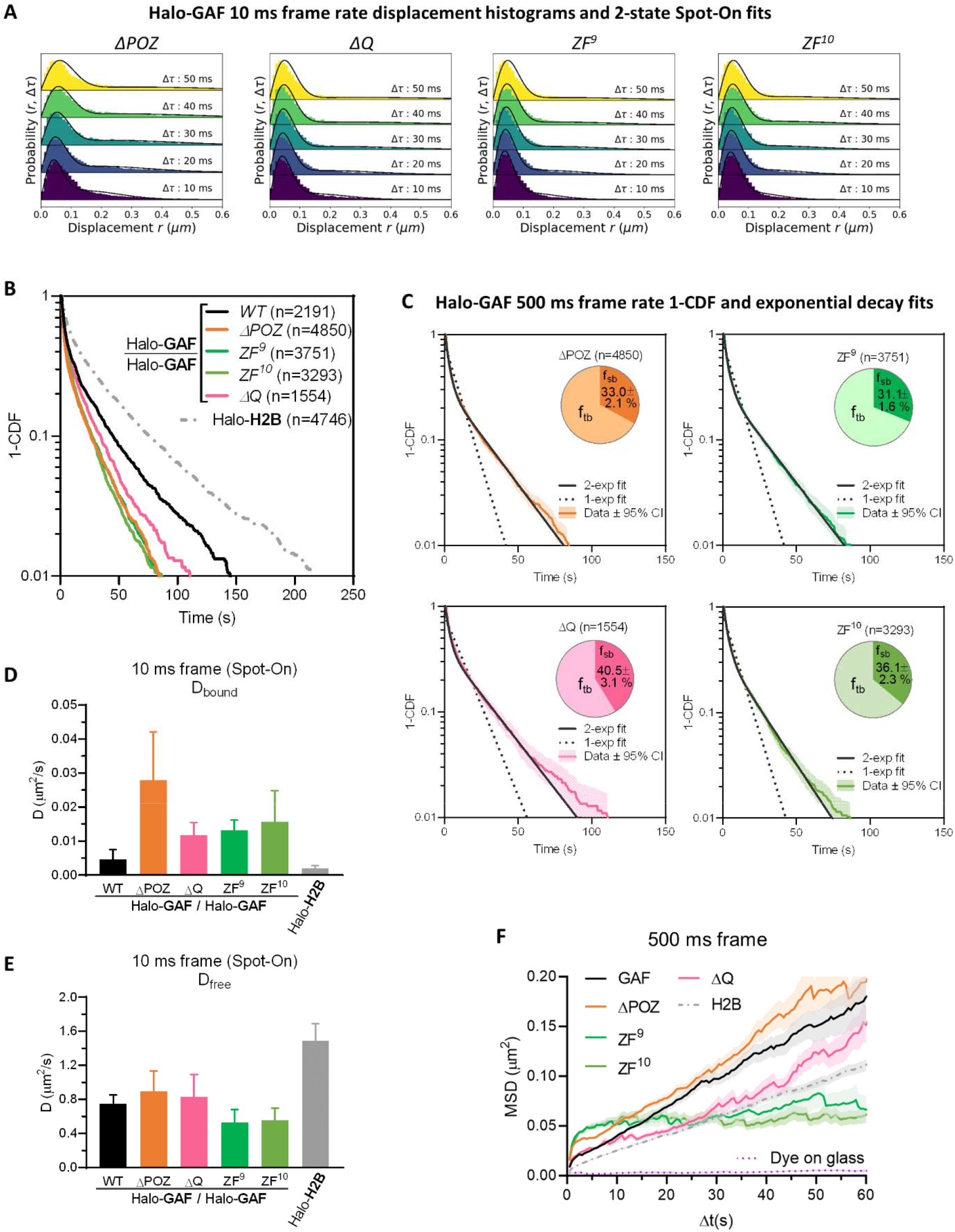
Live-cell SPT diffusive parameters for Halo-GAF mutants. (A) Spot-On fits of fast-tracking data for Halo-GAF mutants (see Fig. S2B for *WT*). (B) Survival probability curves (1-CDF) from apparent dwell times of >1000 single-molecule chromatin-binding events, for *WT* and mutant Halo-GAF. (C) One-component and two-component exponential fit of survival probabilities (1-CDF) from slow tracking data of Halo-GAF mutants (see Fig. S3E for *WT*). Pie charts show the stable-binding (f_sb_) and transient-binding (f_sb_) fractions derived from two-component fits. (D) Diffusion coefficients of bound fraction (D_bound_) for Halo-GAF and Halo-H2B derived by Spot-On. (E) Diffusion coefficients of free fraction (D_free_) for Halo-GAF and Halo-H2B derived by Spot-On. (F) Average MSD versus lag time for *WT* and Halo-GAF mutants at 500 ms frame rate. Mean and SE (shaded) are shown. System noise is shown by the MSD of dye molecules stuck on coverglass.

**Figure S6.**
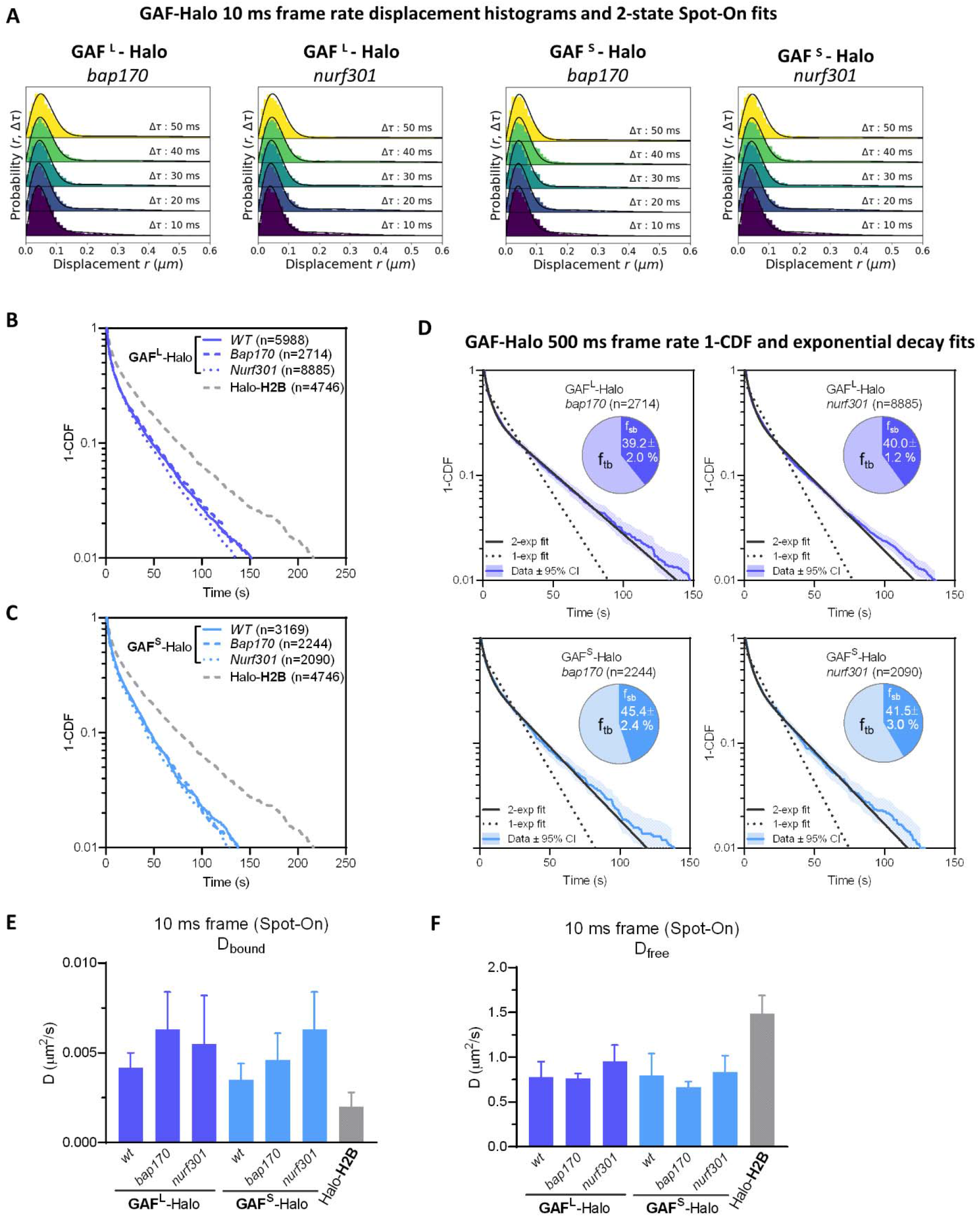
Live-cell SPT diffusive parameters for GAFL-Halo and GAFS-Halo in *bap170* and *nurf301* mutants. (A) Spot-On fits of fast-tracking data for GAF^L^-Halo and GAF^S^-Halo in *bap170* and *nurf301* mutants (see Fig. S2B for *WT*). See methods for genotypes of *WT, bap170*, and *nurf301*. (B) Survival probability curves (1-CDF) from apparent dwell times of more >1000 single-molecule chromatin-binding events for GAF^L^-Halo in *WT, bap170* and *nurf301* mutants. (C) Survival probability curves (1-CDF) from apparent dwell times of more >1000 single-molecule chromatin-binding events for GAF^S^-Halo in *WT, bap170* and *nurf301* mutants. (D) One-component and two-component exponential fit of survival probabilities (1-CDF) from slow tracking data for GAF^L^-Halo and GAF^S^-Halo in *bap170* and *nurf301* mutants (see Fig. S3D for *WT*). Pie charts show the stable-binding (f_sb_) and transient-binding (f_sb_) fractions derived from two-component fits. (E) Diffusion coefficients of bound fraction (D_bound_) for GAF^L^-Halo, GAF^S^-Halo and Halo-H2B derived by Spot-On. (F) Diffusion coefficients of free fraction (D_free_) for GAF^L^-Halo, GAF^S^-Halo and Halo-H2B derived by Spot-On.

**Figure S7.**
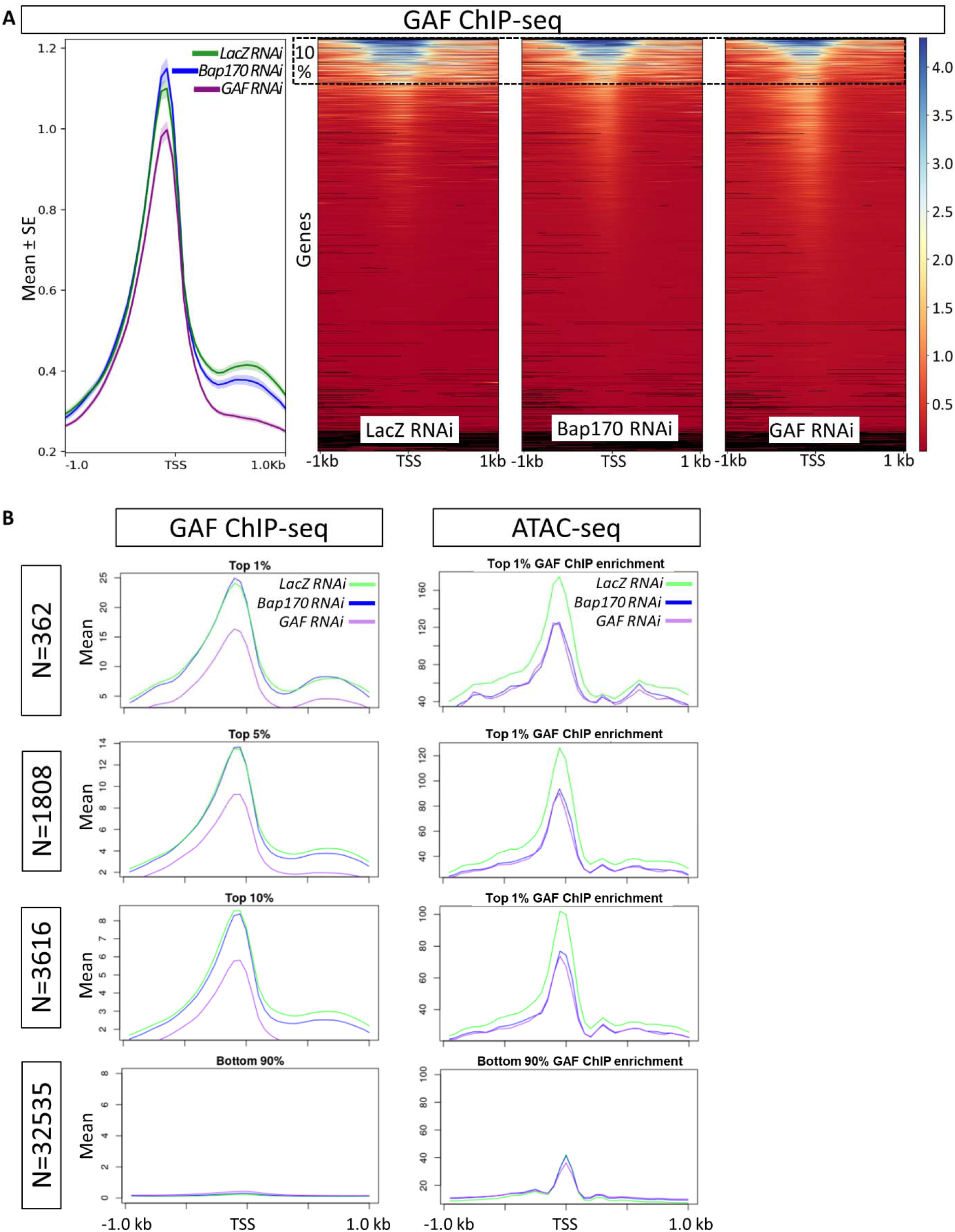
Comparison of GAF ChIP-seq signals in LacZ RNAi (control) and Bap170 RNAi experiments of Judd et al. 2021. (A) Comparison of GAF ChIP-seq signals in LacZ RNAi (control), Bap170 RNAi and GAF RNAi experiments (bw files from GSE157235) derived from^24^. The left graph shows mean ChIP enrichment (mean±SE) for all regions ±1kb centered around transcription start sites (TSS); The right graphs show heat maps of all genes (generated by computeMatrix/plotHeatmap of deepTools^109^). Dashed rectangle indicates the top 10% regions with the highest GAF ChIP enrichment in control (for which the mean enrichment is plotted in (B)). (B) Comparison of GAF ChIP-seq, ATAC-seq, and PRO-seq signals in LacZ RNAi (control), Bap170 RNAi and GAF RNAi experiments (bw files from GSE157235, GSE149336, and GSE149332, respectively) derived from^24^. Regions ±1kb flanking TSS were sorted according to mean GAF ChIP enrichment in LacZ RNAi from high to low as shown in (A). Mean values of GAF ChIP enrichment (left column) and ATAC-seq (right column) enrichment are plotted for the top 1%, 5%, 10% of regions with the highest GAF ChIP signal and for the remaining 90% regions. For 3616 TSS-flanking regions with highest GAF ChIP enrichment, on average, chromatin accessibilities (ATAC-seq) are reduced in both *Bap170 RNAi* and *GAF RNAi* conditions, while the mean enrichment for GAF ChIP-seq shows no change in *Bap170 RNAi*. These analyses indicate that although there are differential effects at specific sites, the overall genome-wide GAF chromatin binding is not affected in PBAP-depleted condition.

**Figure S8.**
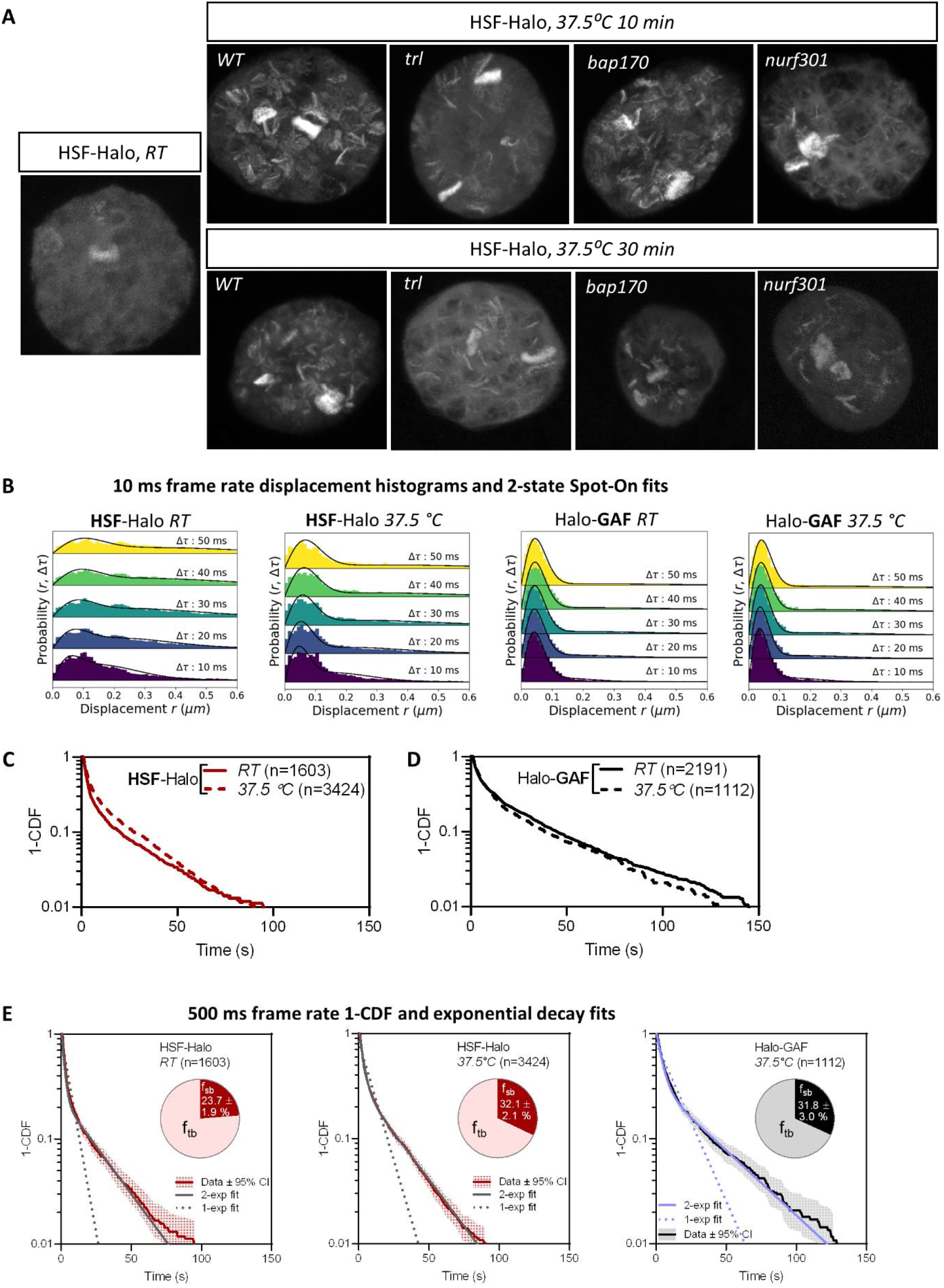
HSF-Halo binding on polytene chromosomes and live-cell SPT at RT and HS conditions. (A) Confocal images of HSF-Halo in fixed salivary glands. HSF-Halo is mostly nucleoplasmic at room temperature (*RT*) and binds to many loci after heat shock (*HS*) at 37.5 °C for 10 and 30 min. Maximum projections of confocal z-stacks are shown because major HSF bands are located in different focal planes. The pattern of HSF binding on heat shock is substantially reduced in *trl* and *nurf301* mutants and partially affected in the *bap170* mutant. Polytene loci showing little or no change of HSF binding in the *trl* mutant is consistent with findings that not all HS genes are GAF-dependent^18^. (These genes presumably require an analogous pioneer factor and attendant remodelers). See methods for genotypes of *WT, trl, bap170*, and *nurf301*. (B) Spot-On fits of fast-tracking data for HSF-Halo (*RT, 37.5°C*) and Halo-GAF (*37.5°C*, see Fig. S2B for *RT*). (C) Survival-probability curves (1-CDF) from apparent dwell times of >1000 single-molecule chromatin-binding events for HSF-Halo at *RT* and *37.5°C*. (D) Survival-probability curves (1-CDF) from apparent dwell times of >1000 single-molecule chromatin-binding events for Halo-GAF at *RT* and *37.5°C*. (E) One-component and two-component exponential fit of survival probabilities (1-CDF) from slow tracking data for HSF-Halo (RT, *37.5°C*) and Halo-GAF (*37.5°C*, see Fig. S3D for *RT*). Pie charts show the stable-binding (f_sb_) and transient-binding (f_sb_) fractions derived from two-component fits.

**Figure S9.**
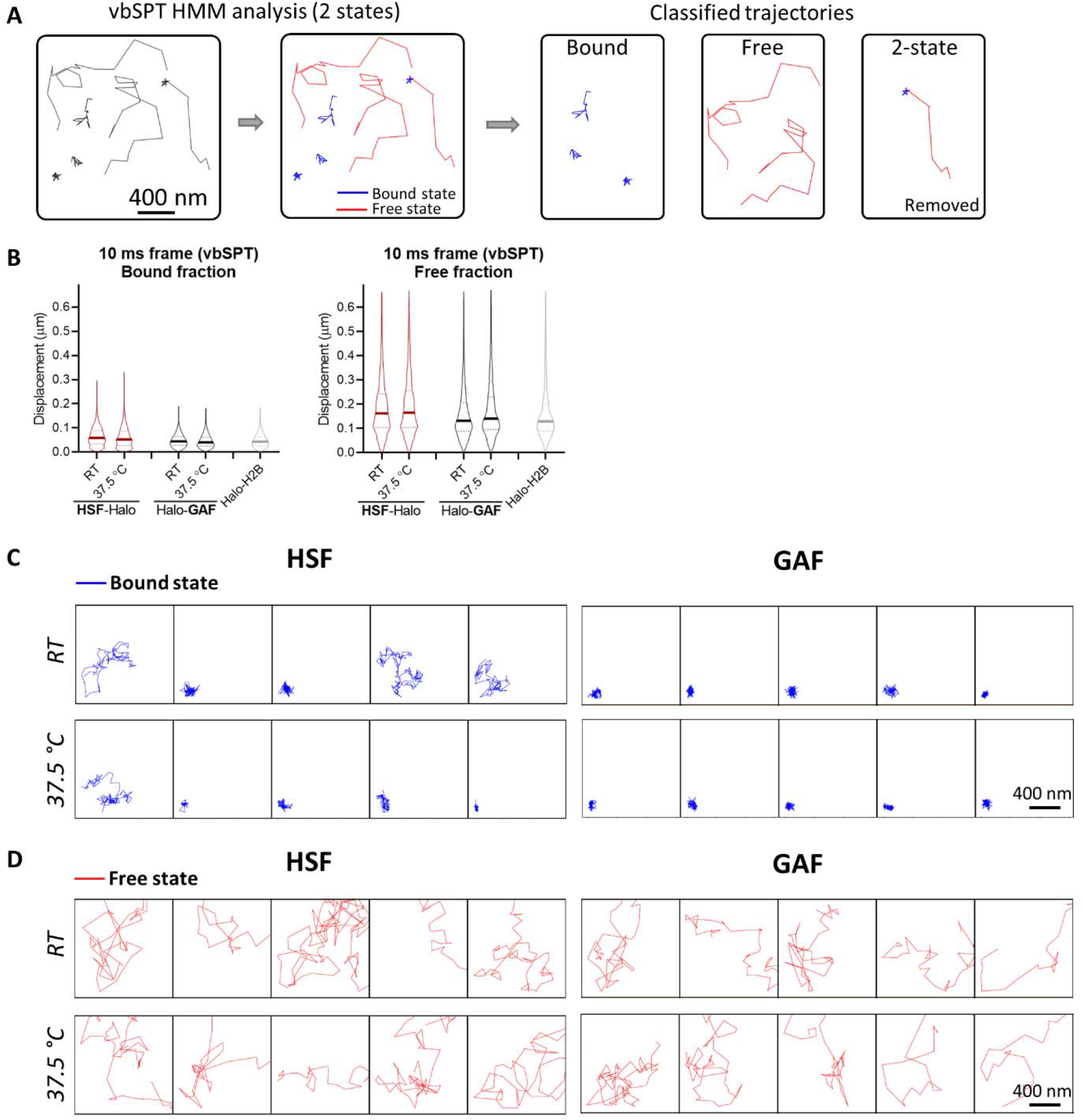
vbSPT analysis of fast-tracking data for HSF-Halo and Halo-GAF at *RT* and *HS* conditions. (A) Overview of fast-tracking trajectory classification with displacement-based HMM classification (vbSPT). After assigning each displacement as either in bound or free state, each trajectory is sub-classified as ‘bound’ or ‘free’, a small fraction of trajectories containing 2 states are excluded from the following analysis in (B-G) and (Fig. 5). (B) Violin plots of displacements show distinct distributions for bound and free trajectories classified by vbSPT. (C) Examples of bound trajectories classified by vbSPT for HSF-Halo, Halo-GAF at *RT* and *37.5°C*, and Halo-H2B at *RT*. (D) Examples of free trajectories classified by vbSPT for HSF-Halo, Halo-GAF at *RT* and *37.5°C*, and Halo-H2B at *RT*.

**Figure S10.**
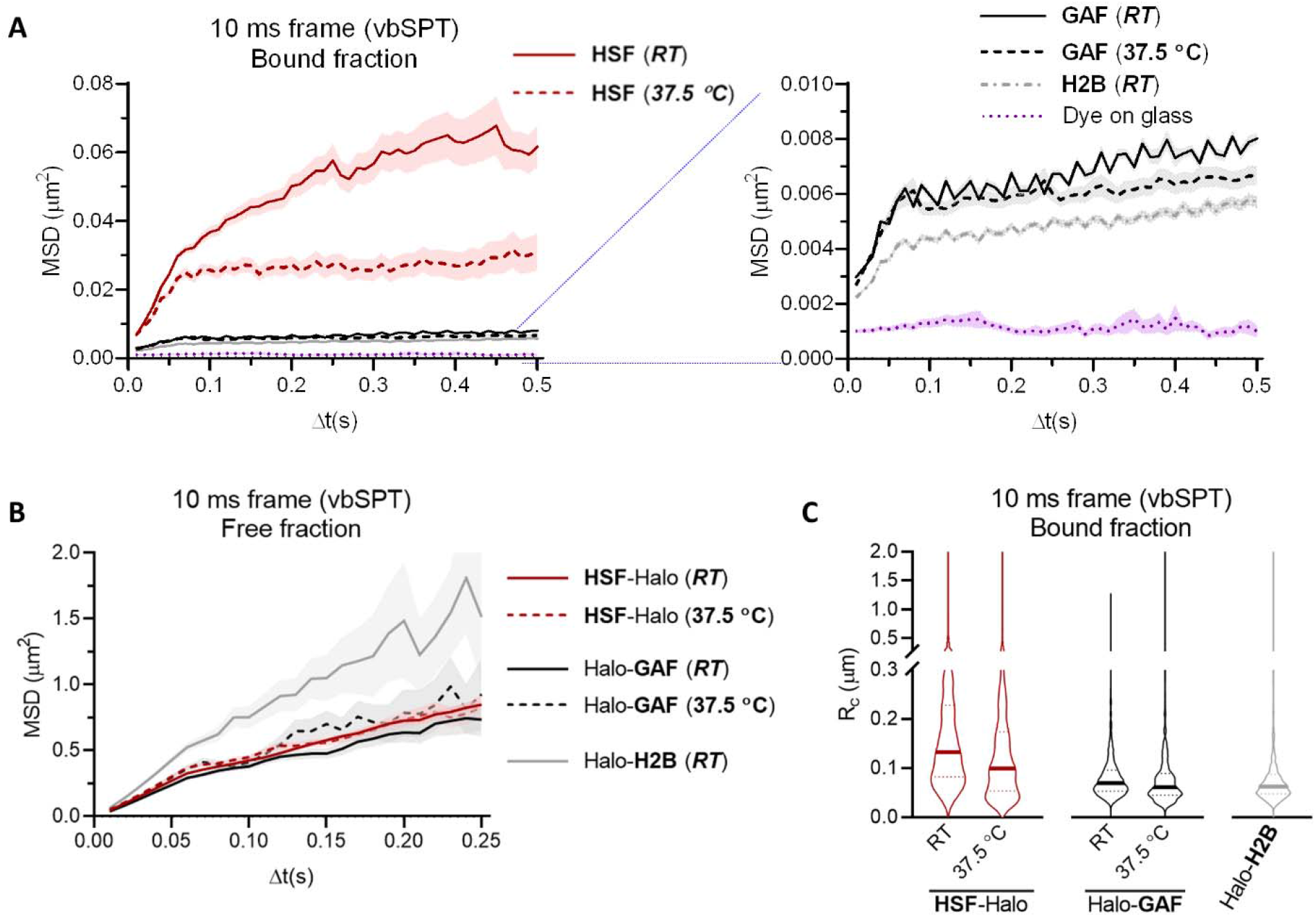
MSD analysis of vbSPT-classified HSF-Halo and Halo-GAF trajectories. (A) Plot of average MSD as a function of lag time Δt of bound trajectories classified by vbSPT for HSF-Halo, Halo-GAF at *RT* and *37.5°C*, and Halo-H2B at *RT*. The right panel shows a zoomed-in section of the same plot. System noise is shown by MSD of dye molecules stuck on the coverglass. Mean and SE (shaded) are shown. (B) Average MSD over Δt of free trajectories classified by vbSPT for HSF-Halo, Halo-GAF at *RT* and *37.5°C*, and Halo-H2B at *RT*. Mean and SE (shaded) are shown. (C) Radius of confinement (Rc) is derived by fitting individual MSD curves with a confined diffusion model, for bound trajectories classified by vbSPT, for HSF-Halo, Halo-GAF at *RT* and *37.5°C*, and Halo-H2B at *RT*.

**Figure S11.**
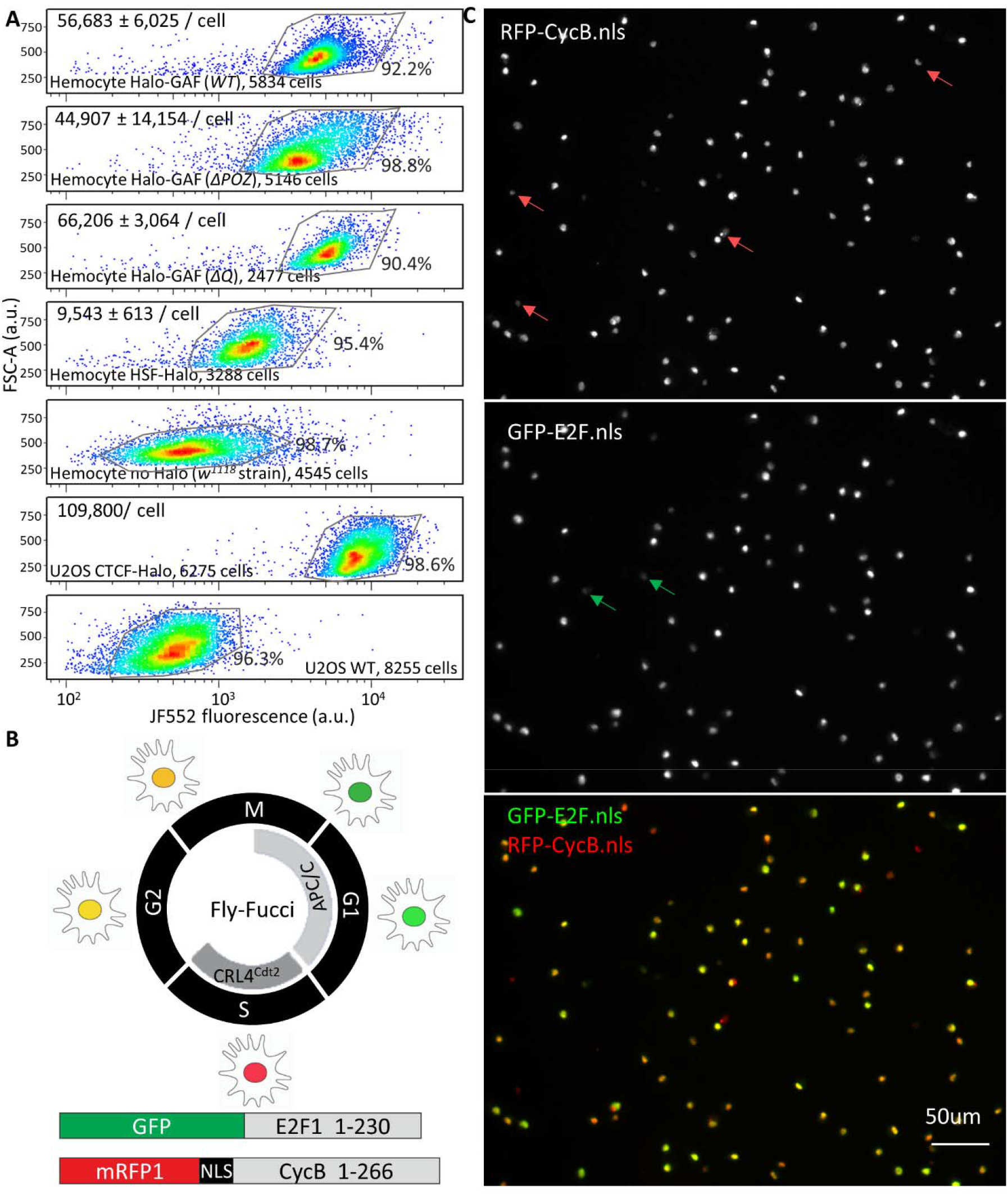
FACS quantitation of Halo-GAF and HSF-Halo in *Drosophila* hemocytes and cell cycle phase identification. (A) Total Halo-GAF (knock-in *WT, ΔPOZ*, and *ΔQ*) and HSF-Halo (transgenic in the *P{PZ}Hsf^03091^/Hsf^3^* background) fluorescence per cell for JF552-stained late 3^rd^ instar larval hemocytes and CTCT-Halo in U2OS cells quantified by flow cytometry. Cellular abundance of Halo-GAF and HSF-Halo molecules are estimated using CTCF-Halo in U2OS cells as a standard (see methods)^81,107^. Hemocytes (*w*^1118^ strain) or U2OS cells not expressing HaloTag were used as controls for background subtraction. One of three representative flow cytometry experiments is shown. Mean ± SD of estimated protein abundance is shown at the upper left corner of each plot. A much larger number of molecules (in the order of one million) for GAF was reported earlier in the S2 cell line^110^; the reason for the discrepancy is unclear. FSC-A, forward scatter area. (B) Conceptual diagram of the Fly-FUCCI system^111^. Both GFP-E2F1_1–230_ and mRFP1-CycB_1–266_ are expressed with the GAL4/UAS system. In early M phase, both GFP-E2F1_1–230_ and mRFP1-CycB_1–266_ are present and thus display yellow. In mid-mitosis, the APC/C marks mRFP1-CycB_1–266_ for proteasomal degradation, leaving the cells fluorescing green. As cells progress from G1 to S phase, CRL4^Cdt2^ degrades GFP-E2F1_1-230_, and cells are labeled red due to the presence of mRFP1-CycB_1–266_ only. After cells enter G2 phase, GFP-E2F1_1-230_ protein levels reaccumulate, marking the cells yellow due to the presence of mRFP1-CycB_1–266_. (C) Characterization of cell-cycle stage for late 3^rd^ instar larval hemocytes. Only 4 out of 96 cells in the field of view show ‘red only’ fluorescence (S phase), and 2 cells have ‘green only’ fluorescence (M to G1 phase). The majority of hemocytes have both red and green fluorescence, indicating G2 phase or early M phase. Given that a previous study shows only 0.32% of larval hemocytes stain positive with the mitotic phosH3 antibody^112^, we conclude that most larval hemocytes are in the G2 phase.

**Movie S1. Fast-tracking movie of Halo-GAF**

A fast-tracking movie of Halo-GAF labeled with 1nM JF554. Movie was acquired with 10 ms camera integration time for single molecule tracking after 10-30 s of initial nuclear glow.

**Movie S2. Slow-tracking movie of GAF^L^-Halo**

A low-tracking movie of GAF^L^-Halo labeled with 0.05 nM JF552 (and 50 nM nonfluorescent JF700 blocker). Images acquired at 500 ms exposure time to motion blur diffusing molecules and selectively observe chromatin-bound molecules. Movie frames are placed on a 3D-axis of time and x, y coordinates to display identified trajectories. Tracking parameters are adjusted to avoid identification of blurred molecules.

