## Supplemental figures for "Kinetic principles underlying pioneer function of GAGA transcription factor in live cells"

**SUPPLEMENTAL FIGURE LEGENDS**

### Figure S1

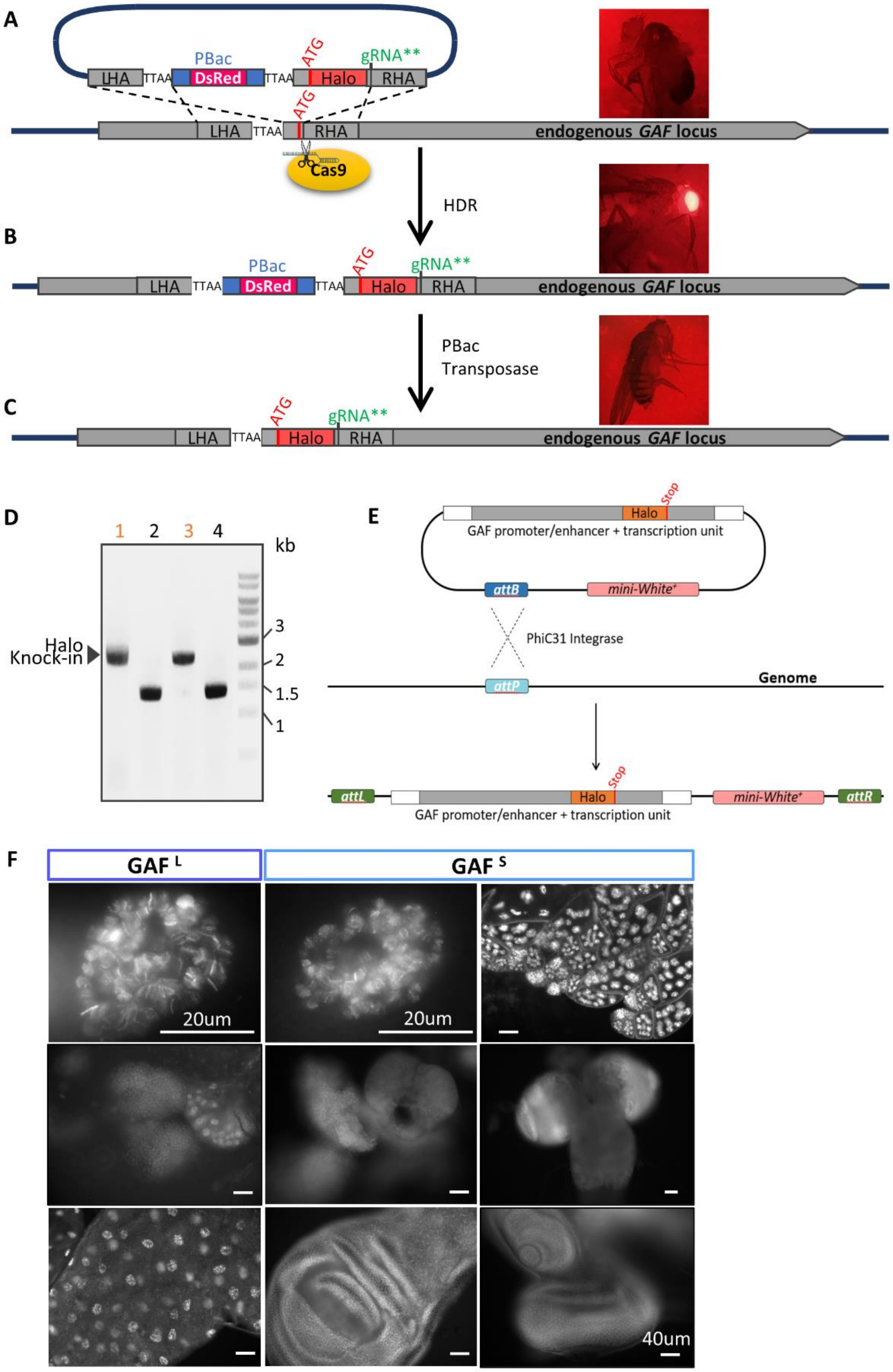

**Figure S1. Generation of N-terminal Halo-GAF knock-in fly strain and C-terminal GAF-Halo transgenic fly strains.**

(A) Donor plasmid design for homology directed repair (HDR). HaloTag and a flexible linker (GGSGS, not shown) are placed downstream of the start codon ATG. A PBac transposon containing a DsRed cassette is inserted into a nearby genomic TTAA site adjacent to the gRNA target site in the coding region that is close to the start codon ATG. The TTAA site is duplicated so that both ends of the PBac transposon contain a TTAA sequence. Approximately 1 kb fragment downstream of the gRNA target site is cloned as the right homology arm (RHA), with silent mutations (gRNA**) introduced to destroy the gRNA PAM sequence in the donor plasmid. Similarly, a 1kb fragment upstream of the genomic TTAA site is cloned as the left homology arm (LHA).

(B) LHA and RHA mediate HDR upon Cas9 cleavage, inserting HaloTag along with the DsRed cassette. Flies that have undergone HDR can be identified by eye DsRed fluorescence.

(C) By crossing to a fly strain expressing PBac transposase, the DsRed cassette can be removed, as indicated by loss of fluorescence, leaving only one TTAA sequence, thereby allowing scarless HaloTag knock-in with a removable selection marker. Arrows indicate positions of the primers used for validating HaloTag insertion.

(D) PCR validation of HaloTag knock-in after DsRed cassette removal (lane 1 and 3). Halo-GAF homozygous flies are viable, showing only 1 band ~900 bp larger than flies without HaloTag knock-in (lane 2 and 4).

(E) Strategy used to generate transgenic fly strains $GAF^{L}$-Halo, $GAF^{S}$-Halo, Halo-H2B and HSF-Halo. An ~15 kb fragment containing the Trl transcription unit and ~1kb upstream and downstream regions was cloned, and HaloTag ORF was inserted upstream of the stop codons for $GAF^{L}$ or $GAF^{S}$, respectively. Thus, each of the two transgenic flies express a Halo-tagged GAFL or GAFS isoform and another isoform (non-tagged), under native Trl promoter control.

(F) Tissue-specific expression of transgenic $GAF^{L}$-Halo and $GAF^{S}$-Halo. Shown are major expressing larval tissues. $GAF^{L}$-Halo: salivary gland, lymph gland, intestine; $GAF^{S}$-Halo: salivary gland, lymph gland, wing disc, ovary, brain, eye-antenna disc.

Figure S2

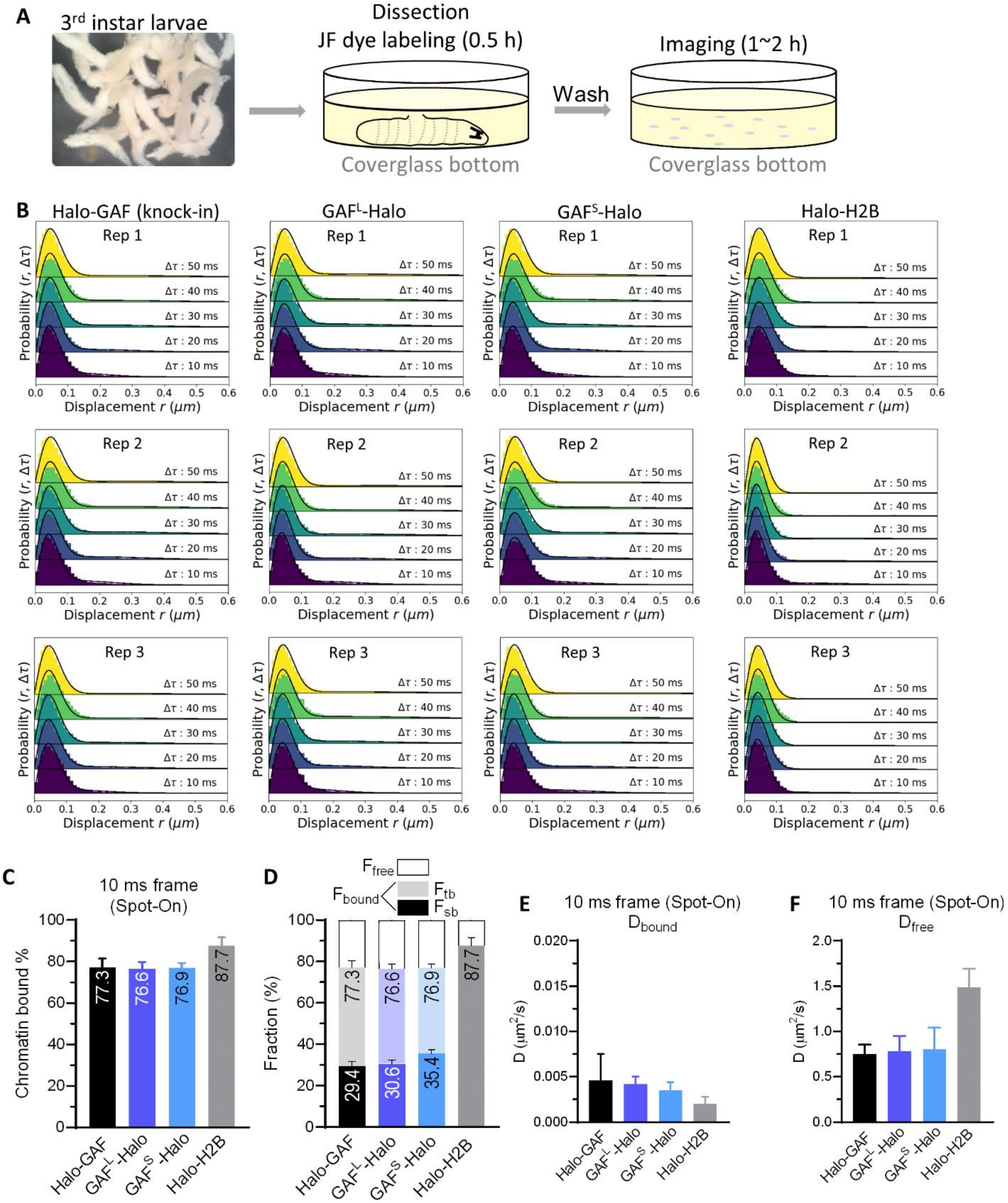


**Figure S2. Hemocyte imaging and fast-tracking diffusive parameters for Halo-GAF, GAF$^{L}$-Halo, GAF$^{S}$-Halo, Halo-H2B.**

(A) Experimental timeline of single-particle imaging with 3rd instar larval hemocytes. 3rd instar larvae are washed with DI H2O (left) and dissected in a coverglass bottom dish containing Schneider's medium and JF dye at room temperature. Upon dissection hemocytes are released into the medium and labeled for 30 min, the rest of the larval tissues are discarded (middle). Cells are briefly washed twice with fresh media and imaged for 1-2 h.

(B) Spot-On fits of Halo-GAF, $GAF^L$-Halo, $GAF^S$-Halo, Halo-H2B fast-tracking data.

(C) Spot-On kinetic modeling of fast-tracking data shows 77% of Halo-GAF is chromatin bound. Similar values are obtained for isoforms $GAF^L$ and $GAF^S$ individually tagged in the presence of untagged GAF isoforms. Results are mean ± SD from three biological replicates.

(D) Chromatin-free fraction ($F_{free}$), long- and short-lived chromatin-binding fractions ($F_{sb}$ and $F_{tb}$) of HaloTagged GAF fusions extracted from fast- and slow-tracking data in (C) and (Fig. S3E), respectively, with error propagation.

(E) Diffusion coefficients of bound fraction ($D_{bound}$) for Halo-GAF, $GAF^L$-Halo, $GAF^S$-Halo, Halo-H2B derived by Spot-On.

(F) Diffusion coefficients of free fraction (Dfree) for Halo-GAF, $GAF^L$-Halo, $GAF^S$-Halo, Halo-H2B derived by Spot-On.

### Figure S3

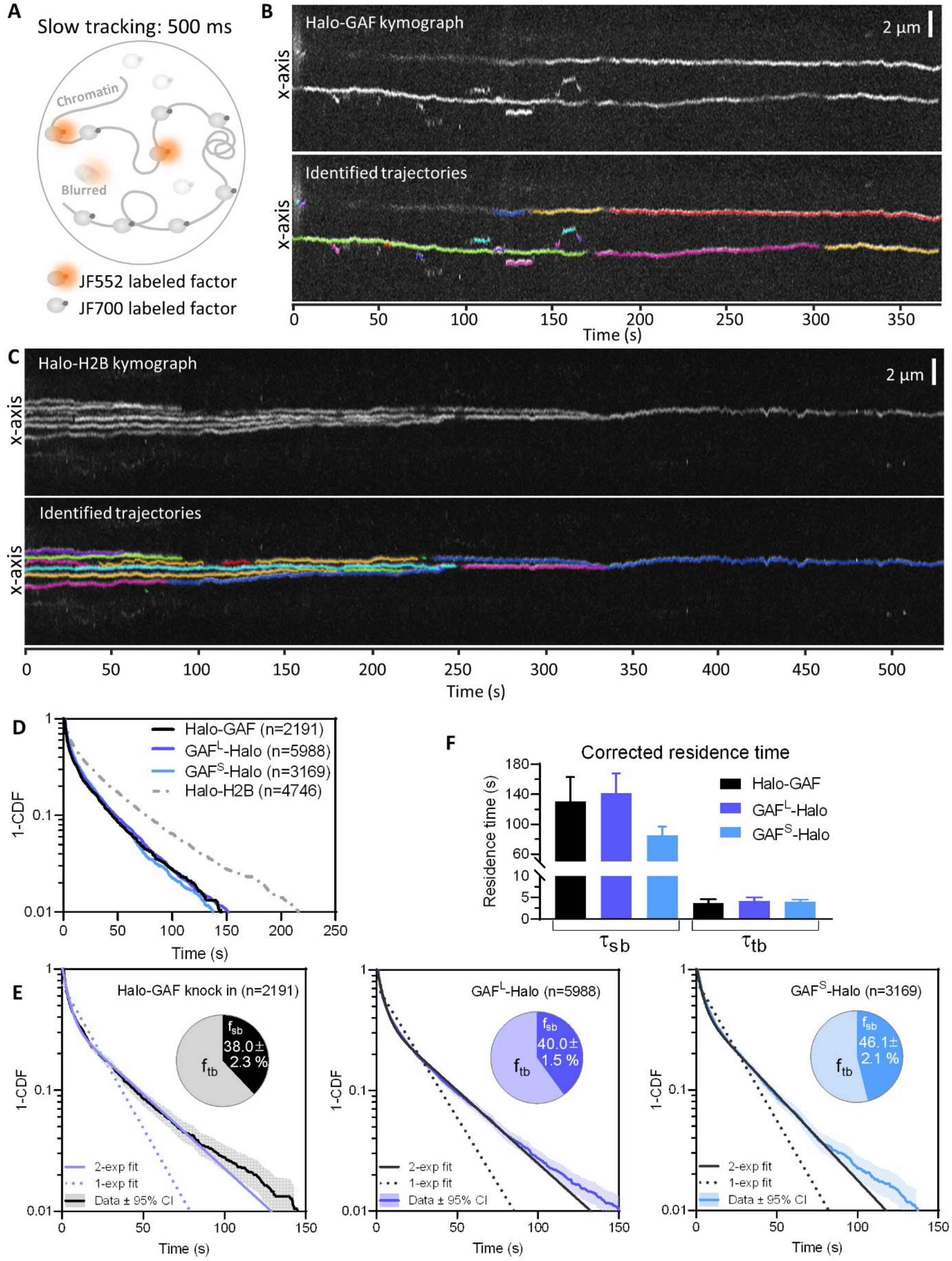


**Figure S3. Slow tracking results for Halo-GAF, GAF$^{L}$-Halo, GAF$^{S}$-Halo, Halo-H2B.**

(A) Fast and slow tracking regimes. Fast tracking with 10 ms frame rate and high laser power allows single molecule imaging to distinguish slow (chromatin-bound) and fast (chromatin-free) diffusing subpopulations. Slow tracking uses low-intensity excitation and 500 ms exposure time to motion blur diffusing molecules and selectively observe the dwell times of chromatin-bound molecules. A higher concentration of JF700 is added to block labelling of most HaloTag protein fusions, while a much lower concentration of JF552 is used to sparsely label a small fraction of HaloTag so that each nucleus shows only 2~10 molecules per frame during image acquisition.

(B) Kymograph of a Halo-GAF slow tracking movie shows traces of bound GAF molecules over time (upper). Trajectories identified from the raw movie are plotted on the kymograph using separate colors (lower).

(C) Kymograph of a Halo-H2B slow tracking movie shows traces of bound H2B molecules over time (upper). Trajectories identified from the raw movie are plotted on the kymograph (lower).

(D) Survival probability curves (1-CDF) plotted from apparent dwell times of thousands (n) of single-molecule chromatin-binding events for Halo-GAF, GAFL-Halo and GAFS-Halo.

(E) One-component and two-component exponential fit of survival probabilities (1-CDF) from slow tracking data of Halo-GAF, $GAF^{L}$-Halo and $GAF^{S}$-Halo. Pie charts show the stable-binding ($f_{sb}$) and transient-binding ($f_{sb}$) fractions derived from two-component fits, and errors represent bootstrapped SD.

(F) Corrected average residence times for stable- ($\tau_{sb}$) and transient- ($\tau_{tb}$) binding by transgenic $GAF^{L}$-Halo and $GAF^{S}$-Halo.

### Figure S4

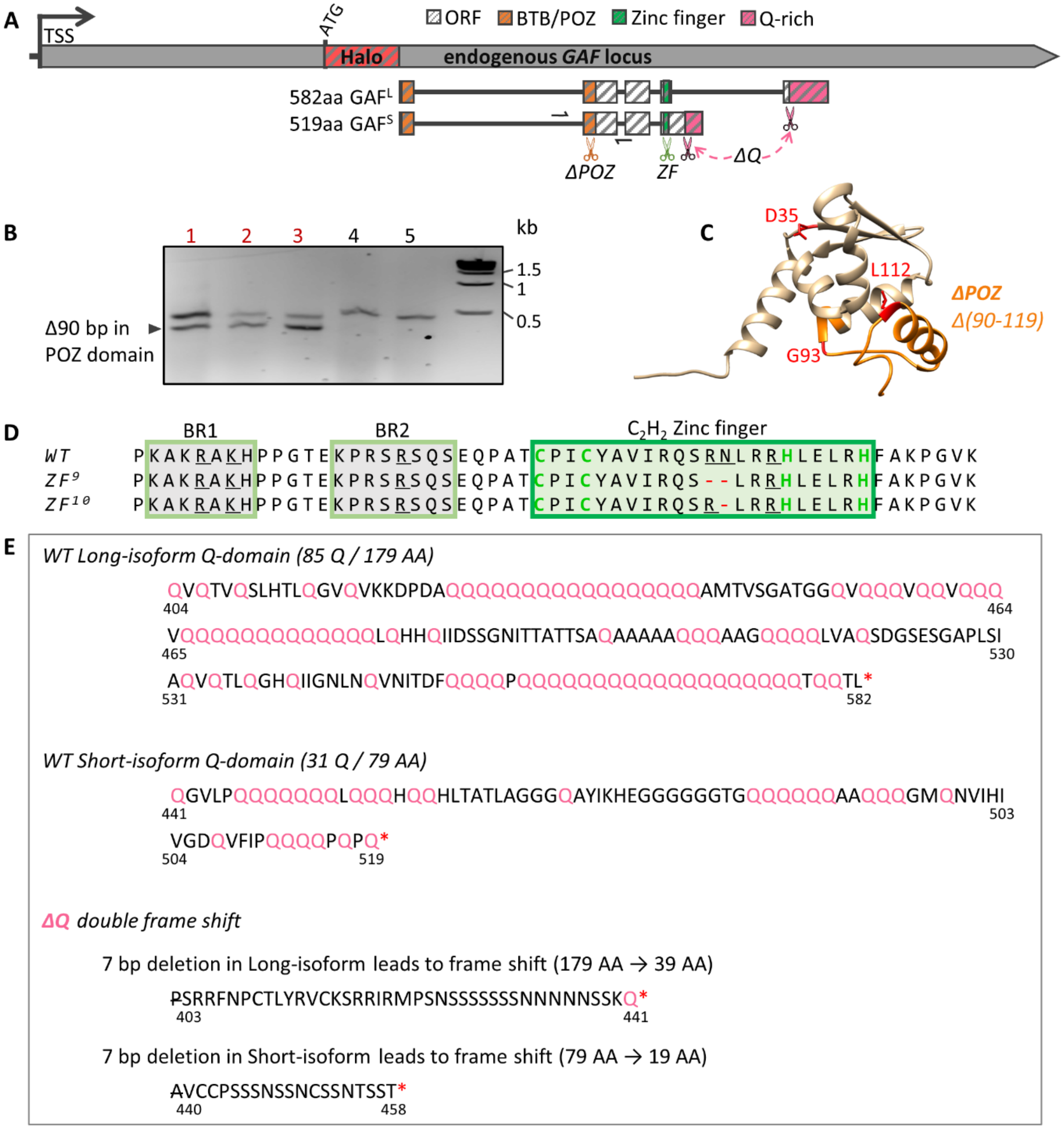


**Figure S4. Generation of mutations in functional domains of Halo-GAF by CRISPR/Cas9 gene editing.**

(A) In the Halo-GAF fly strain, Cas9 and gRNA were introduced to target the BTB/POZ domain, zinc finger, and Q-rich domains, respectively. The BTB/POZ domain is separated by a large intron. A gRNA target site in the second exon (orange scissors) was selected and a donor plasmid containing a 90 bp deletion (*ΔPOZ*) was constructed for homology-directed repair (HDR). For zinc finger mutations, we selected a gRNA target site in the zinc finger coding region (green scissors), and screened for in-frame small deletions generated by non-homologous end joining (NHEJ). To generate deletions of both Q-rich domains in long and short isoforms (*ΔQ*), two gRNAs targeting the upstream ends of two Q-rich domains (pink scissors) were introduced at the same time, and we screened for double frame-shift deletions induced by NHEJ. After gRNA and donor plasmid injection, fly progeny were crossed with balancer chromosome, candidates homozygous-lethal or with reduced viabilities were verified by PCR and sanger sequencing. Half arrows indicate positions of the PCR primers used in (B). TSS, transcription start site.

(B) PCR validation of *ΔPOZ*. Lanes 1-3 show two PCR bands indicating precise deletion in one allele; lanes 4-5 are two lines without the precise deletion. Sanger sequencing verified a precise 90 bp deletion in one allele.

(C) POZ domain model structure. 90bp deletion in the second exon generates a 30-AA deletion (*Δ90-119*) of the POZ domain (*ΔPOZ*, orange), which includes G93 and L112 (red) that have been shown to be essential for transcription activation[29].

(D) Amino acid sequence of GAF DNA binding domain, which contains a single C2H2 zinc finger (green rectangle) and two upstream basic regions (BR1 and BR2, yellow rectangle). Amino acids involved in recognizing the GAGAG consensus sequence are underlined[108]. Two zinc finger mutations were isolated and verified by sanger sequencing, $ZF^{9}$ and $ZF^{10}$, with R356 and N357, or R356 deleted, respectively.

(E) Amino acid sequence of GAF Q-rich domains for long and short isoforms. In *ΔQ*, a 7bp deletion was identified by sanger sequencing in both isoforms at the upstream ends of Q-rich domains, resulting in frameshifts and truncations of the Q-rich domains from both isoforms. P403 in the long isoform and A440 in the short isoform are deleted and the subsequent amino acids are newly introduced by the frame shifts.

### Figure S5

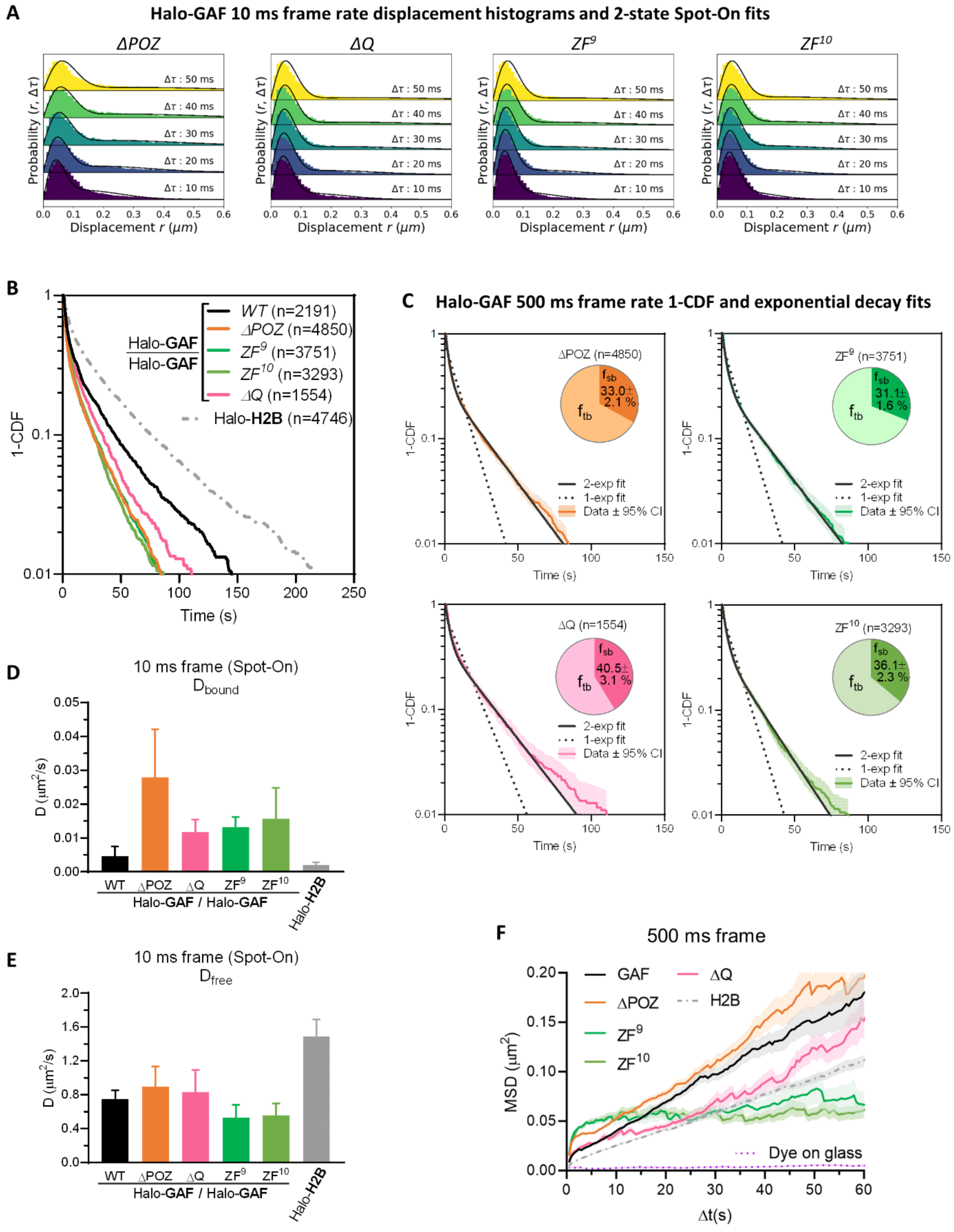


**Figure S5. Live-cell SPT diffusive parameters for Halo-GAF mutants**

(A) Spot-On fits of fast-tracking data for Halo-GAF mutants (see Fig. S2B for *WT*).

(B) Survival probability curves (1-CDF) from apparent dwell times of >1000 single-molecule chromatin-binding events, for *WT* and mutant Halo-GAF.

(C) One-component and two-component exponential fit of survival probabilities (1-CDF) from slow tracking data of Halo-GAF mutants (see Fig. S3E for *WT*). Pie charts show the stable-binding ($f_{sb}$) and transient-binding ($f_{sb}$) fractions derived from two-component fits.

(D) Diffusion coefficients of bound fraction ($D_{bound}$) for Halo-GAF and Halo-H2B derived by Spot-On.

(E) Diffusion coefficients of free fraction ($D_{free}$) for Halo-GAF and Halo-H2B derived by Spot-On.

(F) Average MSD versus lag time for *WT* and Halo-GAF mutants at 500 ms frame rate. Mean and SE (shaded) are shown. System noise is shown by the MSD of dye molecules stuck on coverglass.

### Figure S6

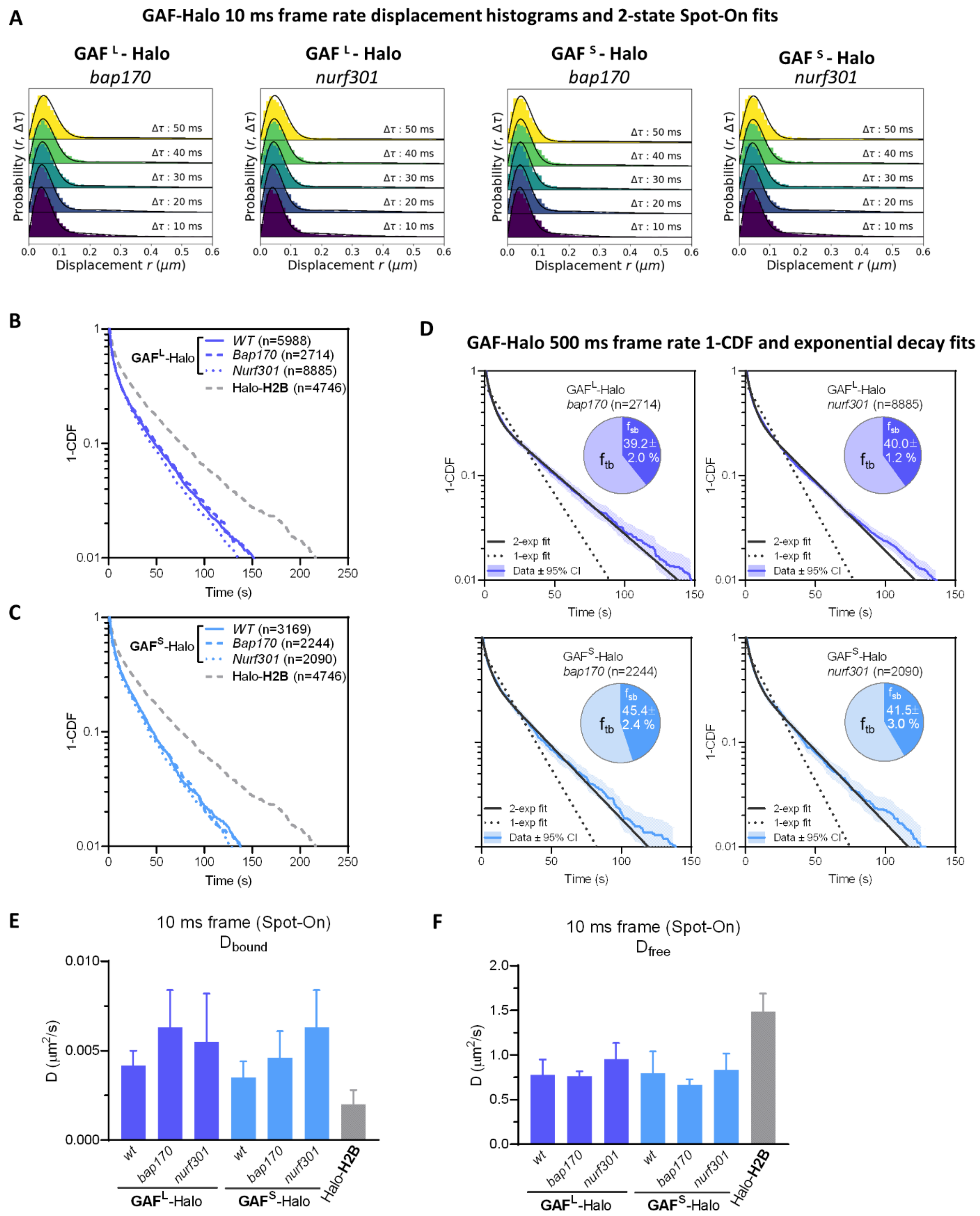


**Figure S6. Live-cell SPT diffusive parameters for GAFL-Halo and GAFS-Halo in *bap170* and *nurf301* mutants**

(A) Spot-On fits of fast-tracking data for $GAF^L$-Halo and $GAF^S$-Halo in *bap170* and *nurf301* mutants (see Fig. S2B for *WT*). See methods for genotypes of *WT*, *bap170*, and *nurf301*.

(B) Survival probability curves (1-CDF) from apparent dwell times of more >1000 single-molecule chromatin-binding events for $GAF^L$-Halo in *WT*, *bap170* and *nurf301* mutants.

(C) Survival probability curves (1-CDF) from apparent dwell times of more >1000 single-molecule chromatin-binding events for $GAF^S$-Halo in *WT*, *bap170* and *nurf301* mutants.

(D) One-component and two-component exponential fit of survival probabilities (1-CDF) from slow tracking data for $GAF^L$-Halo and $GAF^S$-Halo in *bap170* and *nurf301* mutants (see Fig. S3D for *WT*). Pie charts show the stable-binding ($f_{sb}$) and transient-binding ($f_{sb}$) fractions derived from two-component fits.

(E) Diffusion coefficients of bound fraction ($D_{bound}$) for $GAF^L$-Halo, $GAF^S$-Halo and Halo-H2B derived by Spot-On.

(F) Diffusion coefficients of free fraction ($D_{free}$) for $GAF^L$-Halo, $GAF^S$-Halo and Halo-H2B derived by Spot-On.

### Figure S7

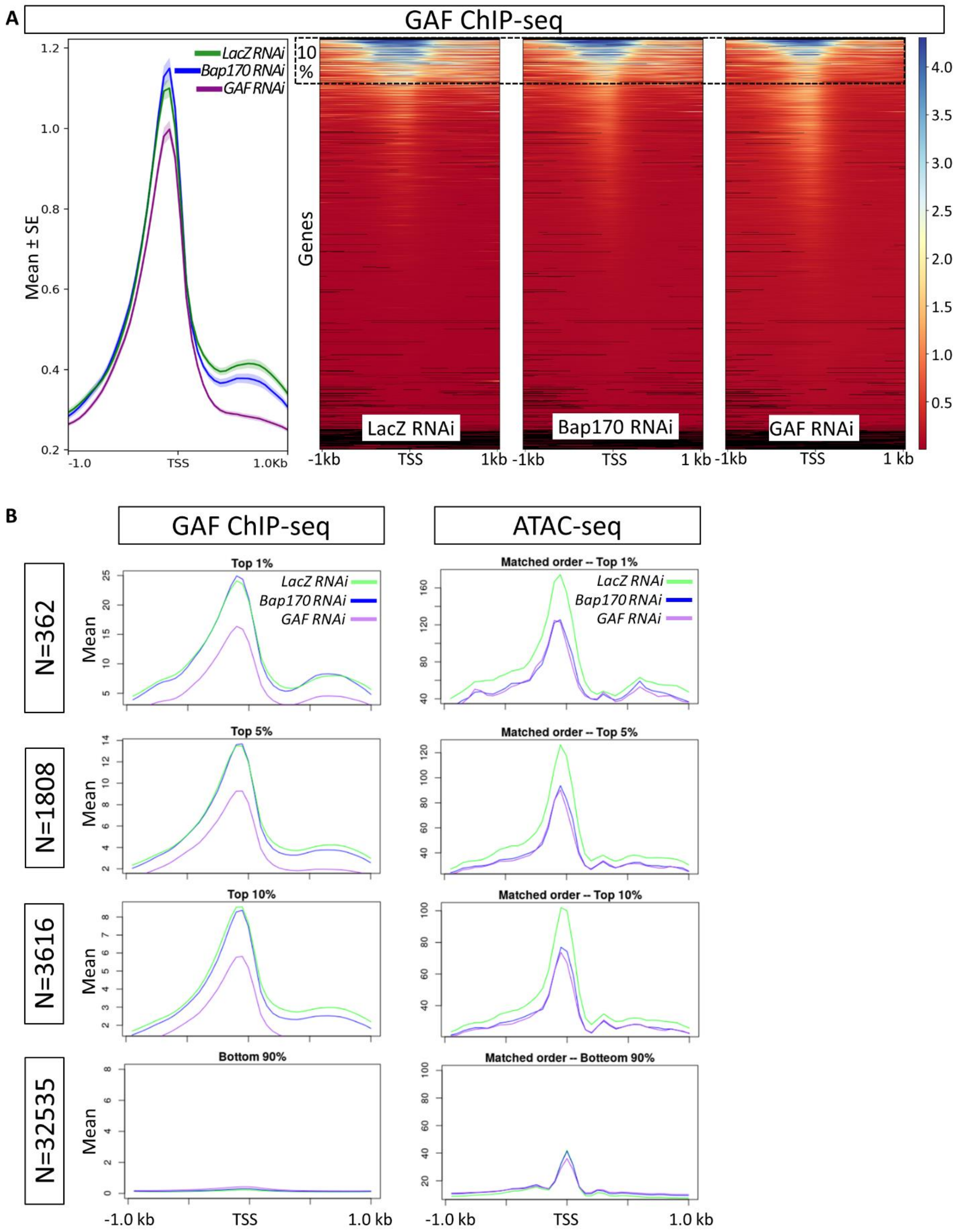

A
GAF ChIP-seq
LacZ RNAi
Bap170 RNAi
GAF RNAi
Mean ± SE
10 %
Genes
LacZ RNAi
Bap170 RNAi
GAF RNAi
-1.0
TSS
1.0Kb
-1kb
TSS
1kb
1 kb
B
GAF ChIP-seq
ATAC-seq
N=362
N=1808
N=3616
N=32535
Top 1%
Top 5%
Top 10%
Bottom 90%
Matched order -- Top 1%
Matched order -- Top 5%
Matched order -- Top 10%
Matched order -- Botteom 90%
Mean
-1.0 kb
TSS
1.0 kb

**Figure S7. Comparison of GAF ChIP-seq signals in LacZ RNAi (control) and Bap170 RNAi experiments of Judd et al. 2021.**

(A) Comparison of GAF ChIP-seq signals in LacZ RNAi (control), Bap170 RNAi and GAF RNAi experiments (bw files from GSE157235) derived from[24]. The left graph shows mean ChIP enrichment (mean±SE) for all regions ±1kb centered around transcription start sites (TSS); The right graphs show heat maps of all genes (generated by computeMatrix/plotHeatmap of deepTools[109]). Dashed rectangle indicates the top 10% regions with the highest GAF ChIP enrichment in control (for which the mean enrichment is plotted in (B)).

(B) Comparison of GAF ChIP-seq, ATAC-seq, and PRO-seq signals in LacZ RNAi (control), Bap170 RNAi and GAF RNAi experiments (bw files from GSE157235, GSE149336, and GSE149332, respectively) derived from[24]. Regions ±1kb flanking TSS were sorted according to mean GAF ChIP enrichment in LacZ RNAi from high to low as shown in (A). Mean values of GAF ChIP enrichment (left column) and ATAC-seq (right column) enrichment are plotted for the top 1%, 5%, 10% of regions with the highest GAF ChIP signal and for the remaining 90% regions. For 3616 TSS-flanking regions with highest GAF ChIP enrichment, on average, chromatin accessibilities (ATAC-seq) are reduced in both *Bap170 RNAi* and *GAF RNAi* conditions, while the mean enrichment for GAF ChIP-seq shows no change in *Bap170 RNAi*. These analyses indicate that although there are differential effects at specific sites, the overall genome-wide GAF chromatin binding is not affected in PBAP-depleted condition.

### Figure S8

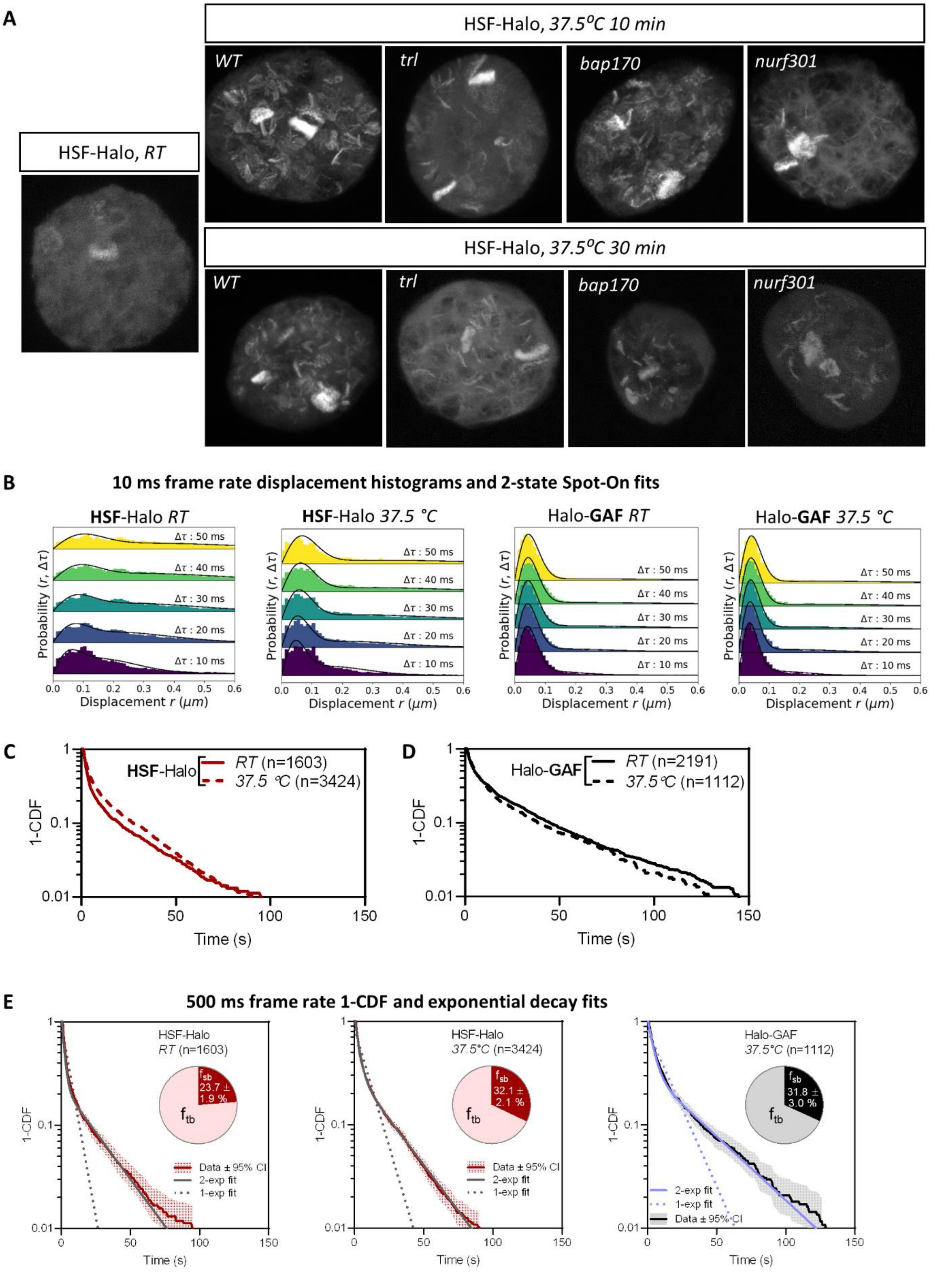
A
HSF-Halo, 37.5°C 10 min
WT
trl
bap170
nurf301
HSF-Halo, RT
HSF-Halo, 37.5°C 30 min
WT
trl
bap170
nurf301
B
10 ms frame rate displacement histograms and 2-state Spot-On fits
HSF-Halo RT
HSF-Halo 37.5 °C
Halo-GAF RT
Halo-GAF 37.5 °C
Δτ : 50 ms
Δτ : 40 ms
Δτ : 30 ms
Δτ : 20 ms
Δτ : 10 ms
Probability (r, Δτ)
Displacement r (μm)
C
HSF-Halo
RT (n=1603)
37.5 °C (n=3424)
1-CDF
Time (s)
D
Halo-GAF
RT (n=2191)
37.5 °C (n=1112)
1-CDF
Time (s)
E
500 ms frame rate 1-CDF and exponential decay fits
HSF-Halo
RT (n=1603)
fsb
23.7 ± 1.9 %
ftb
Data ± 95% CI
2-exp fit
1-exp fit
HSF-Halo
37.5°C (n=3424)
fsb
32.1 ± 2.1 %
ftb
Data ± 95% CI
2-exp fit
1-exp fit
Halo-GAF
37.5°C (n=1112)
fsb
31.8 ± 3.0 %
ftb
2-exp fit
1-exp fit
Data ± 95% CI
1-CDF
Time (s)

**Figure S8. HSF-Halo binding on polytene chromosomes and live-cell SPT at RT and HS conditions.**

(A) Confocal images of HSF-Halo in fixed salivary glands. HSF-Halo is mostly nucleoplasmic at room temperature (*RT*) and binds to many loci after heat shock (*HS*) at 37.5 °C for 10 and 30 min. Maximum projections of confocal z-stacks are shown because major HSF bands are located in different focal planes. The pattern of HSF binding on heat shock is substantially reduced in *trl* and *nurf301* mutants and partially affected in the *bap170* mutant. Polytene loci showing little or no change of HSF binding in the *trl* mutant is consistent with findings that not all HS genes are GAF-dependent[18]. (These genes presumably require an analogous pioneer factor and attendant remodelers). See methods for genotypes of *WT*, *trl*, *bap170*, and *nurf301*.

(B) Spot-On fits of fast-tracking data for HSF-Halo (*RT*, *37.5$^o$C*) and Halo-GAF (*37.5$^o$C*, see Fig. S2B for *RT*).

(C) Survival-probability curves (1-CDF) from apparent dwell times of >1000 single-molecule chromatin-binding events for HSF-Halo at *RT* and *37.5°C*.

(D) Survival-probability curves (1-CDF) from apparent dwell times of >1000 single-molecule chromatin-binding events for Halo-GAF at *RT* and *37.5°C*.

(E) One-component and two-component exponential fit of survival probabilities (1-CDF) from slow tracking data for HSF-Halo (RT, *37.5$^o$C*) and Halo-GAF (*37.5$^o$C*, see Fig. S3D for *RT*). Pie charts show the stable-binding ($f_{sb}$) and transient-binding ($f_{sb}$) fractions derived from two-component fits.

### Figure S9

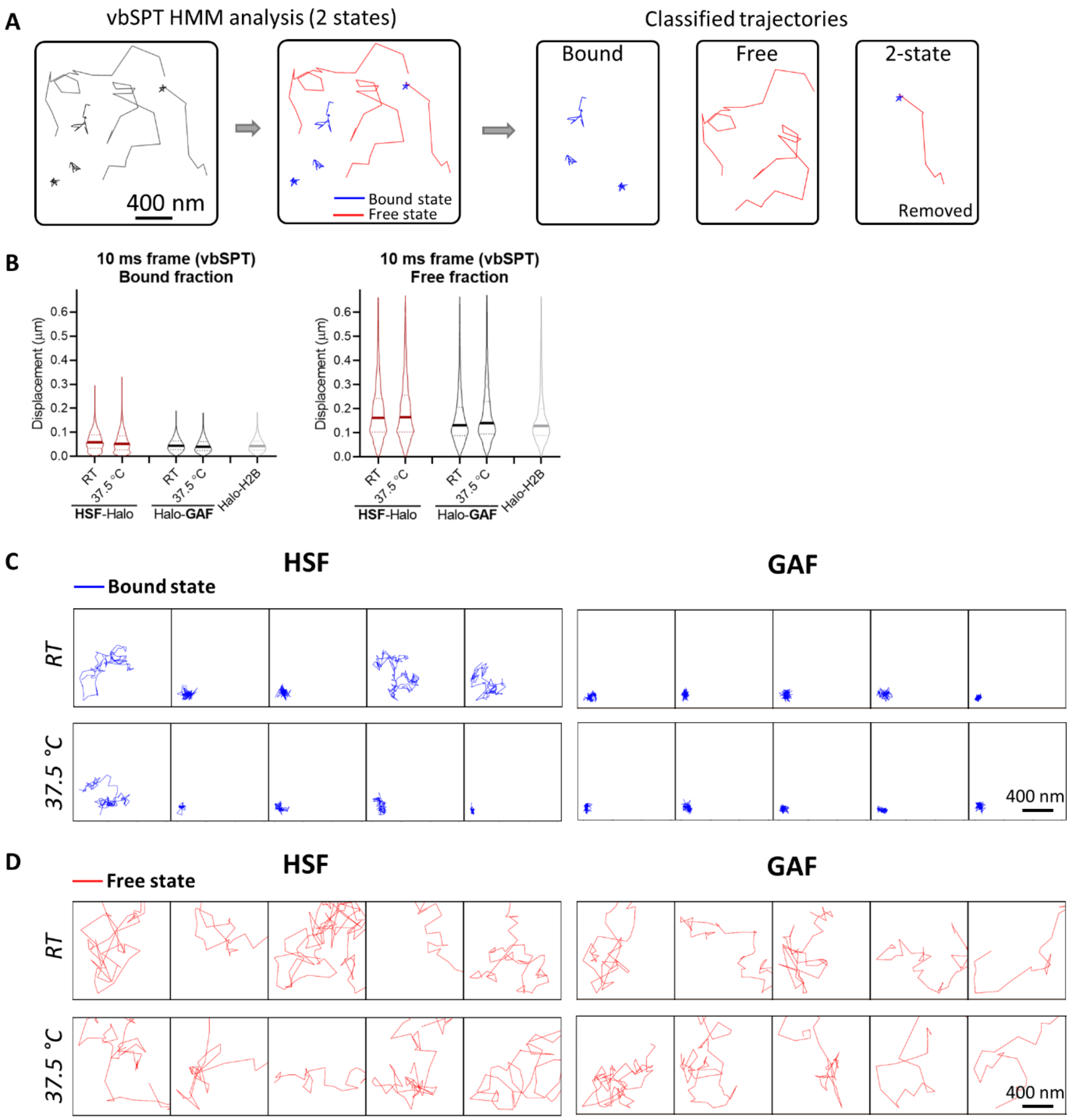


**Figure S9. vbSPT analysis of fast-tracking data for HSF-Halo and Halo-GAF at *RT* and *HS* conditions.**

(A) Overview of fast-tracking trajectory classification with displacement-based HMM classification (vbSPT). After assigning each displacement as either in bound or free state, each trajectory is sub-classified as 'bound' or 'free', a small fraction of trajectories containing 2 states are excluded from the following analysis in (B-G) and (Fig. 5).

(B) Violin plots of displacements show distinct distributions for bound and free trajectories classified by vbSPT.
(C) Examples of bound trajectories classified by vbSPT for HSF-Halo, Halo-GAF at *RT* and *37.5°C*, and Halo-H2B at *RT*.
(D) Examples of free trajectories classified by vbSPT for HSF-Halo, Halo-GAF at *RT* and *37.5°C*, and Halo-H2B at *RT*.

### Figure S10

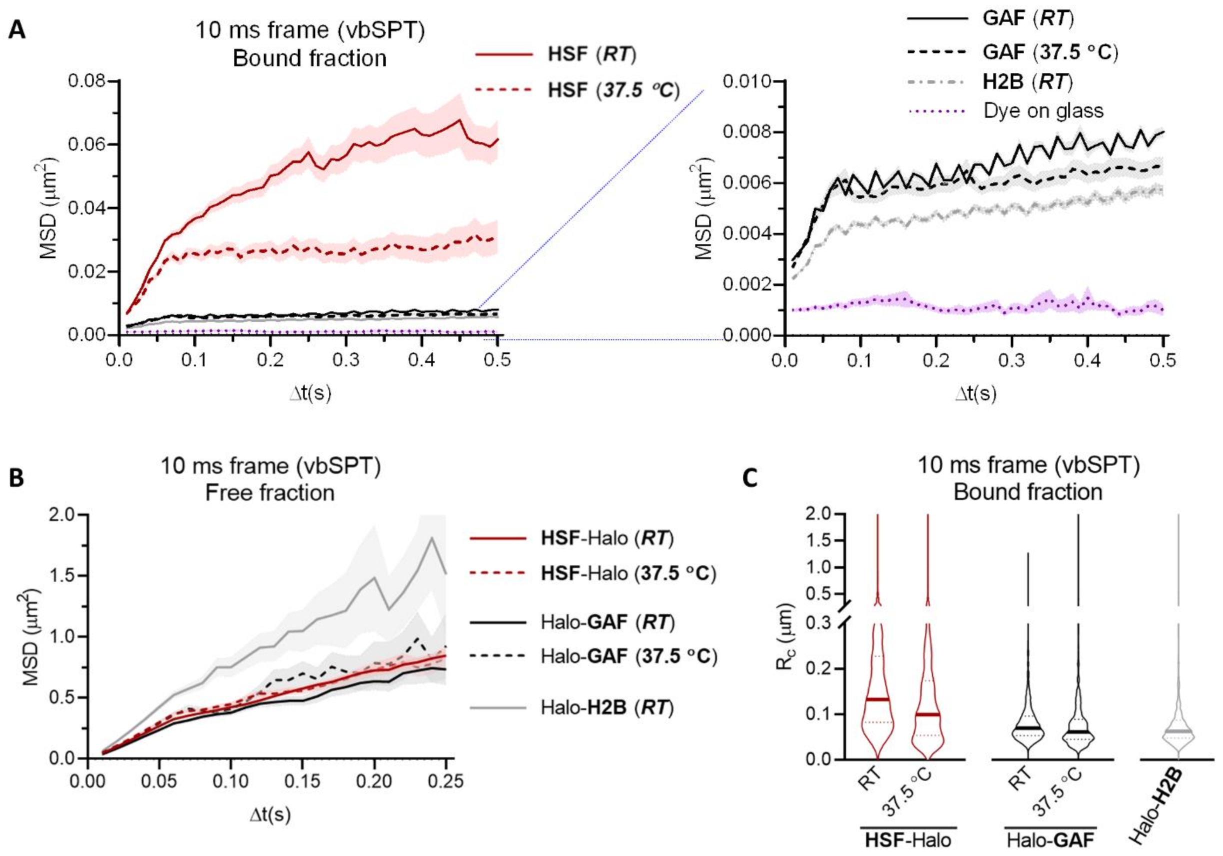


**Figure S10. MSD analysis of vbSPT-classified HSF-Halo and Halo-GAF trajectories.**

(A) Plot of average MSD as a function of lag time Δt of bound trajectories classified by vbSPT for HSF-Halo, Halo-GAF at *RT* and *37.5°C*, and Halo-H2B at *RT*. The right panel shows a zoomed-in section of the same plot. System noise is shown by MSD of dye molecules stuck on the coverglass.
(B) Average MSD over Δt of free trajectories classified by vbSPT for HSF-Halo, Halo-GAF at *RT* and *37.5°C*, and Halo-H2B at *RT*.
(C) Radius of confinement (Rc) is derived by fitting individual MSD curves with a confined diffusion model, for bound trajectories classified by vbSPT, for HSF-Halo, Halo-GAF at *RT* and *37.5°C*, and Halo-H2B at *RT*.

### Figure S11

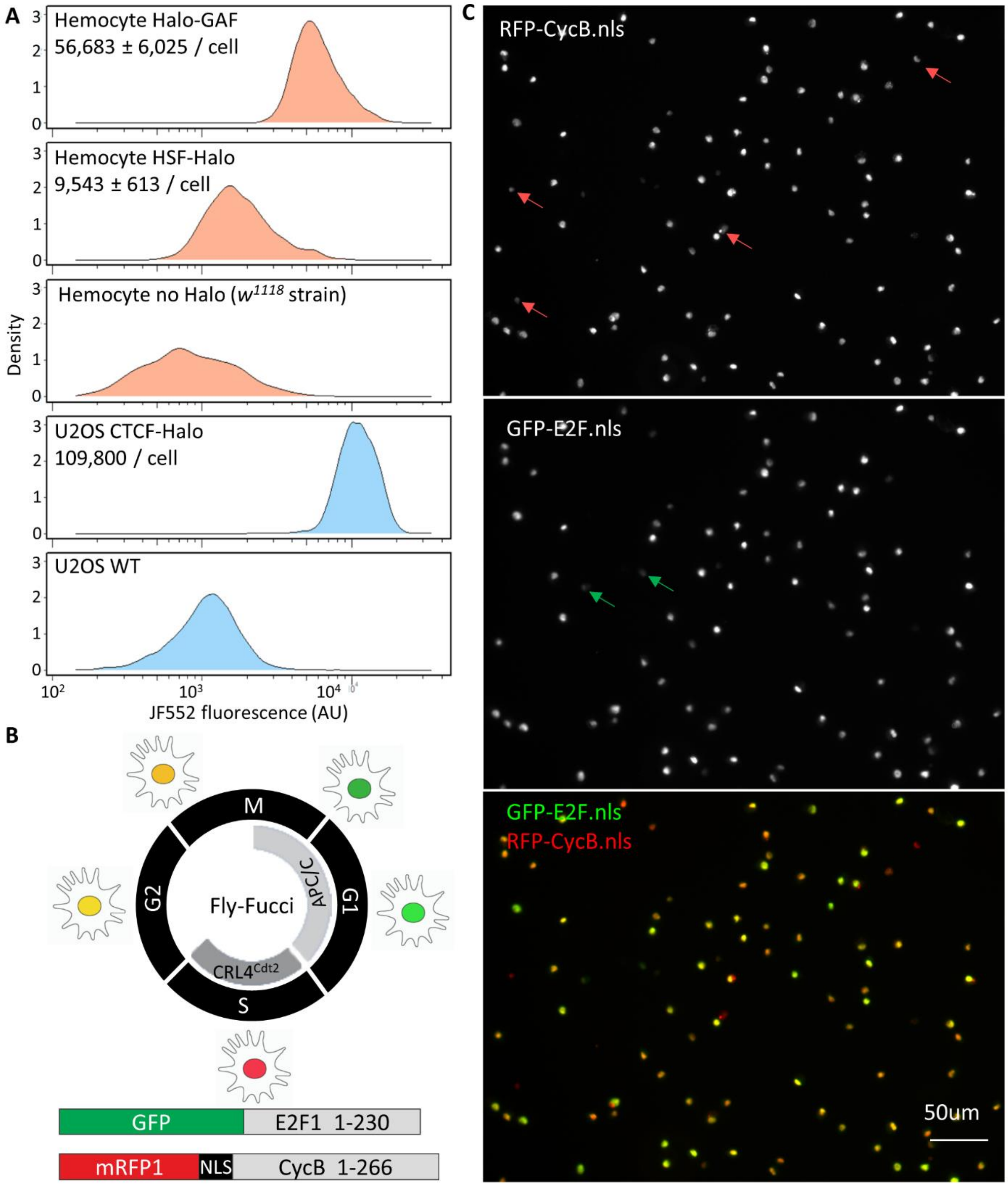


**Figure S11. FACS quantitation of Halo-GAF and HSF-Halo in Drosophila hemocytes and cell cycle phase identification.**

(A) Total Halo-GAF (knock-in) and HSF-Halo (transgenic in the *P{PZ}Hsf*$^{03091}$/*Hsf*$^{3}$ background) fluorescence per cell for JF552-stained late 3$^{rd}$ instar larval hemocytes and calibrated CTCT-Halo in U2OS cells quantified by flow cytometry. Cellular abundance of Halo-GAF and HSF-Halo molecules are estimated using CTCF-Halo in U2OS cells as a standard[81,107]. Hemocytes (*w*$^{1118}$ strain) or U2OS cells not expressing HaloTag were used as controls for background subtraction. Mean ± SD of estimated protein abundance is shown. A much larger number of molecules (in the order of one million) for GAF was reported earlier in the S2 cell line[110]; the reason for the discrepancy is unclear.

(B) Conceptual diagram of the Fly-FUCCI system[111]. Both GFP-E2F1$_{1–230}$ and mRFP1-CycB$_{1–266}$ are expressed with the GAL4/UAS system. In early M phase, both GFP-E2F1$_{1–230}$ and mRFP1-CycB$_{1–266}$ are present and thus display yellow. In mid-mitosis, the APC/C marks mRFP1-CycB$_{1–266}$ for proteasomal degradation, leaving the cells fluorescing green. As cells progress from G1 to S phase, CRL4$^{Cdt2}$ degrades GFP-E2F1$_{1–230}$, and cells are labeled red due to the presence of mRFP1-CycB$_{1–266}$ only. After cells enter G2 phase, GFP-E2F1$_{1–230}$ protein levels reaccumulate, marking the cells yellow due to the presence of mRFP1-CycB$_{1–266}$.

(C) Characterization of cell-cycle stage for late 3$^{rd}$ instar larval hemocytes. Only 4 out of 96 cells in the field of view show ‘red only’ fluorescence (S phase), and 2 cells have ‘green only’ fluorescence (M to G1 phase). The majority of hemocytes have both red and green fluorescence, indicating G2 phase or early M phase. Given that a previous study shows only 0.32% of larval hemocytes stain positive with the mitotic phosH3 antibody[112], we conclude that most larval hemocytes are in the G2 phase.

**Movie S1. Fast-tracking movie of Halo-GAF**

A fast-tracking movie of Halo-GAF labeled with 1nM JF554. Movie was acquired with 10 ms camera integration time for single molecule tracking after 10-30 s of initial nuclear glow.

**Movie S2. Slow-tracking movie of GAF$^{L}$-Halo**

A low-tracking movie of GAF$^{L}$-Halo labeled with 0.05 nM JF552 (and 50 nM nonfluorescent JF700 blocker). Images acquired at 500 ms exposure time to motion blur diffusing molecules and selectively observe chromatin-bound molecules. Movie frames are placed on a 3D-axis of time and x, y coordinates to display identified trajectories. Tracking parameters are adjusted to avoid identification of blurred molecules.
